# Alterations in bile acid metabolizing gut microbiota and specific bile acid genes as a precision medicine to subclassify NAFLD

**DOI:** 10.1101/2020.08.14.251876

**Authors:** Na Jiao, Rohit Loomba, Zi-Huan Yang, Dingfeng Wu, Sa Fang, Richele Bettencourt, Ping Lan, Ruixin Zhu, Lixin Zhu

## Abstract

**Background & Aims:** Multiple mechanisms for the gut microbiome contributing to the pathogenesis of non-alcoholic fatty liver disease (NAFLD) have been implicated. Here, we aim to investigate the contribution and potential application for altered bile acid (BA) metabolizing microbe in NAFLD using whole metagenome sequencing (WMS) data.

**Methods:** 86 well-characterized biopsy-proven NAFLD patients and 38 healthy controls were included in the discovery cohort. Assembly-based analysis was performed to identify BA-metabolizing microbes. Statistical tests, feature selection and microbial interaction analysis were integrated to identify microbial alterations and markers in NAFLD. An independent validation cohort was subjected to similar analyses.

**Results:** NAFLD microbiota exhibited decreased diversity and microbial interactions. We established a classifier model with 53 differential species exhibiting a robust diagnostic accuracy (AUC=0.97) for dectecting NAFLD. Next, 8 important differential pathway markers including secondary BA biosynthesis were identified. Specifically, increased abundance of 7α-HSDH, baiA and baiB were detected in NAFLD. Further, 10 of 50 BA-metabolizing metagenome-assembled genomes (MAG)s, from *Bacteroides ovatus* and *Eubacterium biforme*, were dominant in NAFLD and interplayed as a synergetic ecological guild. Importantly, two subtypes of NAFLD patients were observed according to secondary BA metabolism potentials. Elevated capability for secondary BA biosynthesis was also observed in the validation cohort.

**Conclusions:** We identified novel bacterial BA-metabolizing genes and microbes that may contribute to NAFLD pathogenesis and serve as disease markers. Microbial differences in BA-metabolism and strain-specific differences among patients highlight the potential for precision medicine in NAFLD treatment.

## Introduction

Non-alcoholic fatty liver disease(NAFLD) has become one of the leading causes of liver disease worldwide, with the global prevalence estimated to be 24%.[1] NAFLD is expected to be the No. 1 cause for cirrhosis in the United States within a decade.[2]

The pathogenic mechanism of NAFLD remains unclear. The current multiple-hit hypothesis is that NAFLD is a consequence of a myriad of factors acting in a parallel and synergistic manner in individuals with genetic predisposition.[3] Factors such as insulin resistance, central obesity, environmental or nutritional factors, and gut microbiota, as well as genetic and epigenetic factors, are linked to its pathogenesis.[2, 4, 5]

Recently, the crosstalk between the gut and the liver is increasingly recognized, and many studies have reported dysregulated gut microbiota in NAFLD patients. [6-10] There are several potential mechanisms for the gut microbiota to influence NAFLD development. These effects are mediated by microbial components and metabolites, such as lipopolysaccharide, alcohol, and bile acid(BA).[11]

BA not only facilitate the digestion and absorption of fatty foods as detergent, they also act as important signaling molecules via nuclear receptors, such as farnesoid X receptor(FXR) and G protein coupled BA receptor(GPBAR1 or TGR5) to modulate hepatic BA synthesis, glucose and lipid metabolism. Recently, we observed suppressed BA-mediated FXR signaling in NAFLD liver and intestine, which is in harmony with increased secondary BA production. Furthermore, using 16S rRNA data, we observed elevated abundance of secondary BA metabolizing related bacteria and pathways in the gut microbiome of NAFLD. [12] However, the 16S rRNA sequencing data has limited resolution which does not allow the identification of the species or an accurate functional analysis. [13]

Whole metagenome sequencing(WMS) allows us to achieve a satisfactory resolution of the microbiome. Earlier we have used the WMS data to characterize the gut microbiota in NAFLD patients with and without advanced fibrosis and identified 37 differential bacterial species, among which the abundance of *Escherichia coli* and *Bacteroides vulgatus* was increased in patients with advanced fibrosis and it’s association with microbial metabolites.[9, 14-16] WMS data were also used to study the interactions between the gut microbiome and steatosis in obesity.[15, 17] However, a similar study is lacking for the comparison of the gut community between healthy and NAFLD subjects using WMS data, which is our goal in this study. Here we report the structural and functional characteristics of the gut microbiome in NAFLD, and its association with BA metabolism.

## Results

### Gut microbiota alterations between NAFLD patients and healthy controls

WMS data from 86 well-characterized biopsy-proven NAFLD patients and 38 healthy controls with similar characteristics (Table 1 and Table S1) were chosen to study the structural and functional differences in gut microbiota between NAFLD patients and healthy controls. And we have confirmed that gender or age distribution did not account for the observed microbial differences in this study (Figure S1).

**Table1.** Characteristics of the cohort included in this study.

|  | Discovery cohort |  | Validation cohort |  |
| --- | --- | --- | --- | --- |
|  | NAFLD | Control | NAFLD | Control |
| Sample Size | 86 | 38 | 10 | 11 |
| Age | 51.56±12.67 | 55.71±12.75 | 53.7±3.65 | 56.18±6.65 |
| BMI | 30.25±5.46 | 23.03±1.88 | 34.1±1.2 | 23.19±0.92 |
| Gender(F%/M%) | 44.19/55.81 | 50.00/50.00 | 20.00/80.00 | 63.63/36.36 |
| AST(U/L) | 32.5±29.96 | NA<sup>\$</sup> | 30.8±2.4 | NA |
| LDL cholesterol(mg/dL) | 116±37.12 | NA | 52.25±5.41<sup>#</sup> | NA |
| HDL cholesterol (mg/dL) | 46±15.97 | NA | 20.36±1.26 | NA |
| Triglycerides(mg/dL) | 129±95.70 | NA | 50.45±7.21 | NA |
| Total cholesterol(mg/dL) | 191.5±43.39 | NA | 95.90±5.41 | NA |

Data are presented as median±SD
\$ NA, not available. The control groups included healthy individuals (Ref 33 and 36)

#### Compositional changes in NAFLD gut microbiota

We determined the microbial compositions of NAFLD and healthy controls using WMS data. Bacteroidetes, Firmicutes, Actinobacteria and Proteobacteria were the dominant phyla that collectively account for around 90% proportions in both groups (Figure S2A). NAFLD individuals had lower bacterial diversity than healthy controls (Figure S2B). Besides, significant compositional differences were observed between these two groups (Figure S2C).

To identify microbial markers that may distinguish NAFLD from healthy subjects, differential species were determined with Mann-Whitney U-tests. 53 species with FDR values < 0.1 were identified as differential species (Figure 1 & Table S2). Among these, 11 species were dominant in NAFLD patients, which mainly belong to Clostridia class, including E*ubacterium siraeum, Clostridium bolteae*, E. *coli* and *B.ovatus, B.stercoris* from Bacteroidia class. On the other hand, 42 species significantly reduced in NAFLD patients were mainly of Bacteroidia class, including, Bacteroides dorei, Alistipes shahii, and of Clostridia class, for instance, Eubacterium eligens, Eubacterium hallii, and Faecalibacterium prausnitzii. In addition, random forest (RF) model constructed with differential species achieved an AUC of 0.97 to detect NAFLD patients from controls (Figure S3).

**Figure 1.**
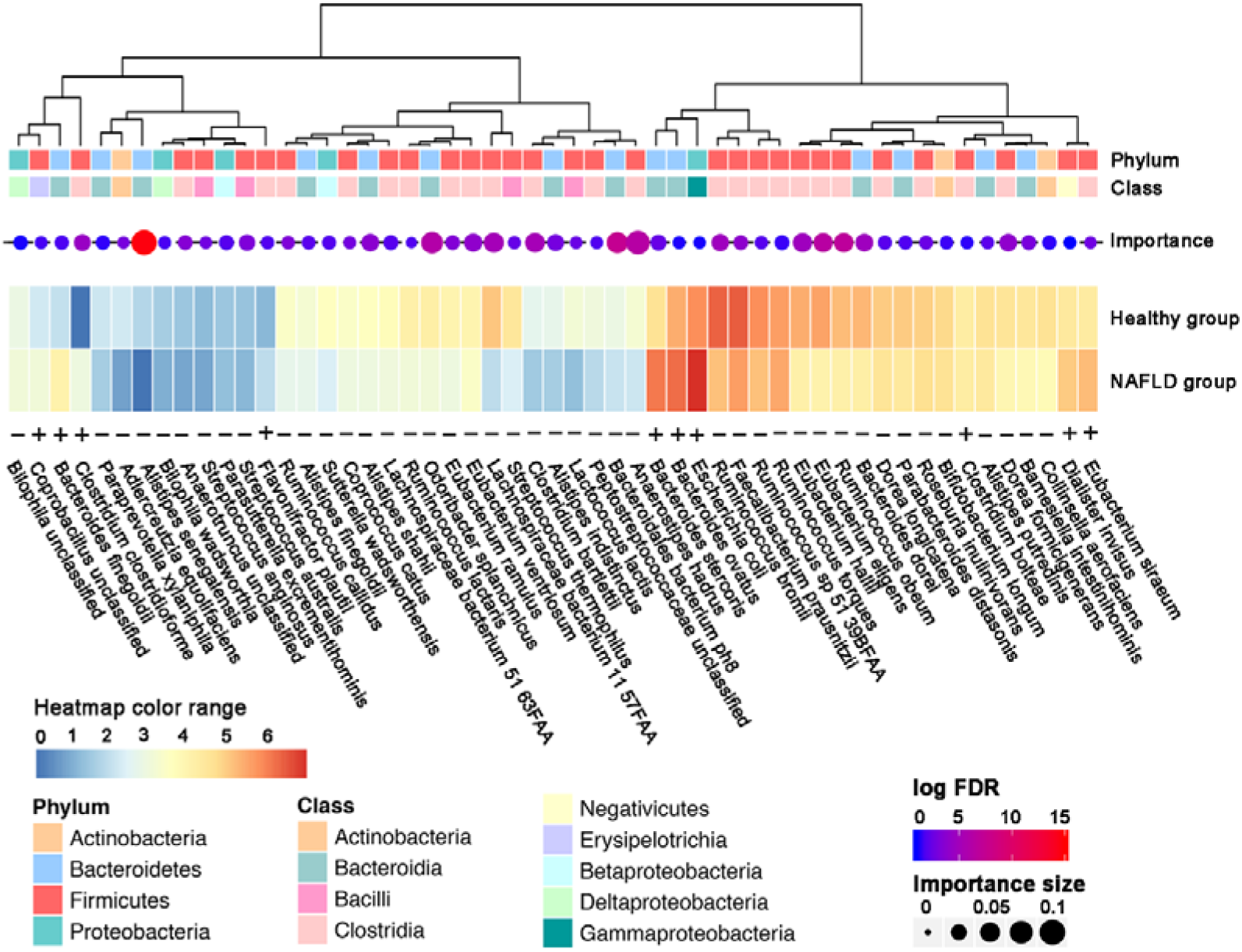
The differential species distinguishing NAFLD patients from healthy controls. Differential species were selected by statistical tests (two-tailed Mann-Whitney U-tests adjusted by Benjamini–Hochberg). Furthermore, the importance of the species that distinguish NAFLD patients from healthy controls was evaluated with random forest model. The heatmap shows the relative abundance (log-transformed) of the differential species in the NAFLD and the healthy groups, the size of the dots is proportional to the importance and the color shows the FDR value (-log-transformed). “+” indicates increased abundance while “-” indicates decreased abundance in NAFLD.

#### Ecological structural changes in NAFLD gut microbiota

Furthermore, at whole-community level, microbial interaction analysis was performed to investigate potential changes in ecological structure. There were more species in healthy communities than those in NAFLD communities (167 nodes vs 141 nodes) though with similar amount of interactions. Then, we examined the “core community” (interactions with magnitudes > 0.4) of healthy and NAFLD groups, respectively. Considerable discrepancies existed in the “core community” of healthy and NAFLD (Figure 2A&B). In detail, the healthy “core community” was more complex, with 162 species and 565 interactions, compared to the NAFLD community with 81 species and 166 interactions. And the NAFLD community was separated into 8 isolated components, an indication of unstable microbial community. Among them, the major component harbored most species from Clostridia class, such as BA production bacteria, *C.bolteae* (node NO. 78), *C.clostridioforme* (node NO. 138) with increased proportion in NAFLD, while species from Bacilli class were dominant in the second major component. Besides, species with increased abundance in NAFLD patients (circle nodes in Figure 2B) were dominant in the “core community” and positively interacted with each other. Then, we looked into the top 20 hub species of “core community”, respectively. 10 of them were common in both group, such as *C.bolteae, C.hathewayi, Dorea longicatena, Flavonifractor plautii*, which may play the role as the “keystone” to sustain the homeostasis (Figure 2C&D).

**Figure 2.**
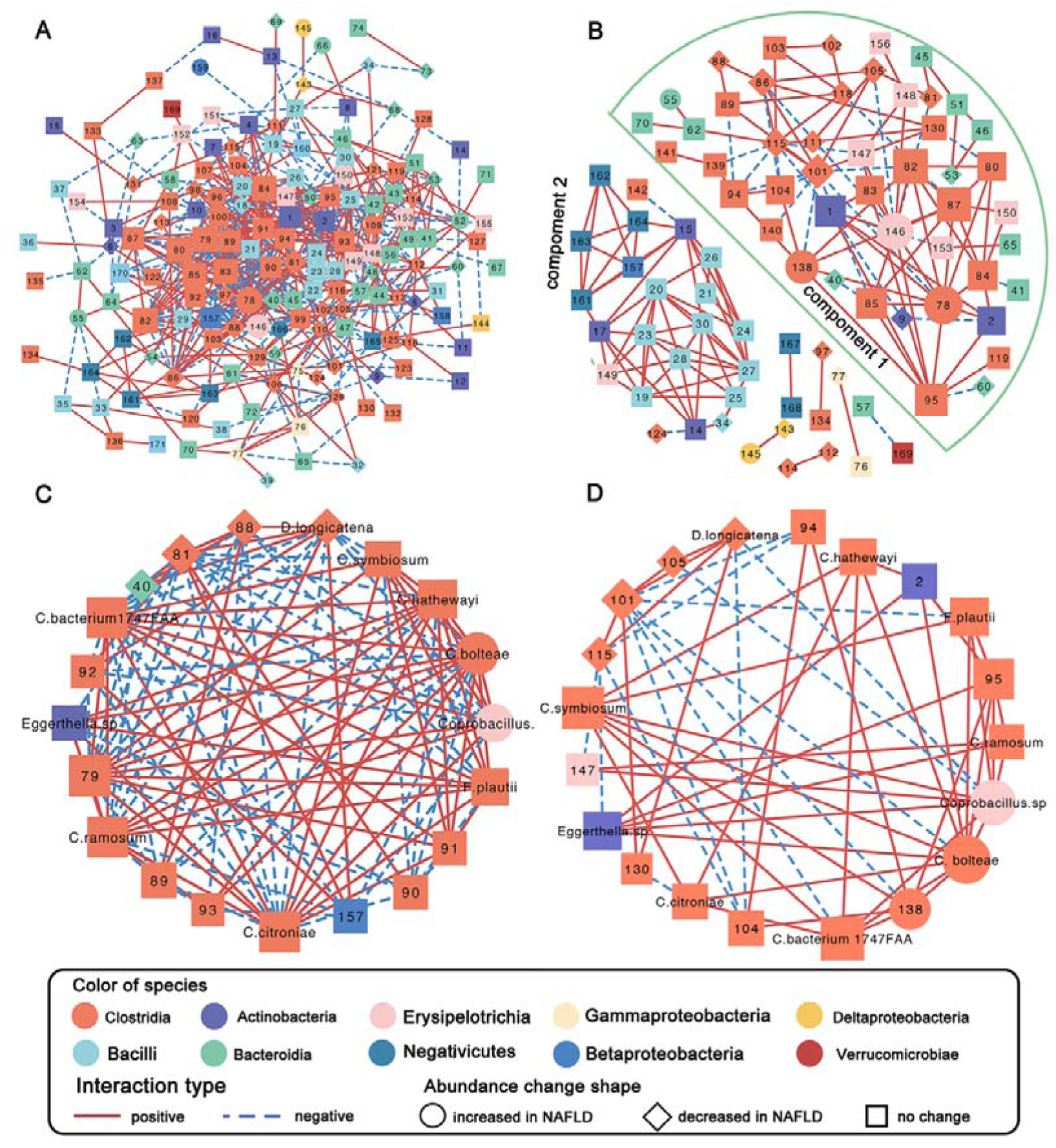
Microbiota “core community” in healthy controls (A&C) and NAFLD patients (B&D). The microbial interactions were calculated using SparCC with 100 refining interactions, and p value of each interaction is approximated with 1000 permutations. Only interactions with p value < 0.05 and interactions with magnitudes > 0.4 were included in the “core community”. The species were colored according to the class they belong to and the node size indicates the hub score in their community. Sub-network of top 20 hub nodes in healthy community (C) and NAFLD community (D) was also plotted. The nodes indicated by species name were common species in both sub-networks.

#### Functional changes in NAFLD gut microbiota

Microbial functional profiles were determined at pathway level using HUMAnN2 and 92 differential pathways were identified between the NAFLD and the healthy groups (Table S3). Similarly, we identified 8 important pathway features (Figure 3A) to build RF model (AUC=0.83) that could distinguish NAFLD patients from healthy subjects (Figure 3B). Most pathways were more represented in NAFLD microbiota than in controls. These pathways included secondary BA synthesis (ko00121) (Figure 3C), benzoate degradation (ko00362), biosynthesis of ansamycins (ko01051) and oxidative phosphorylation (ko00190) (Figure S4).

**Figure 3.**
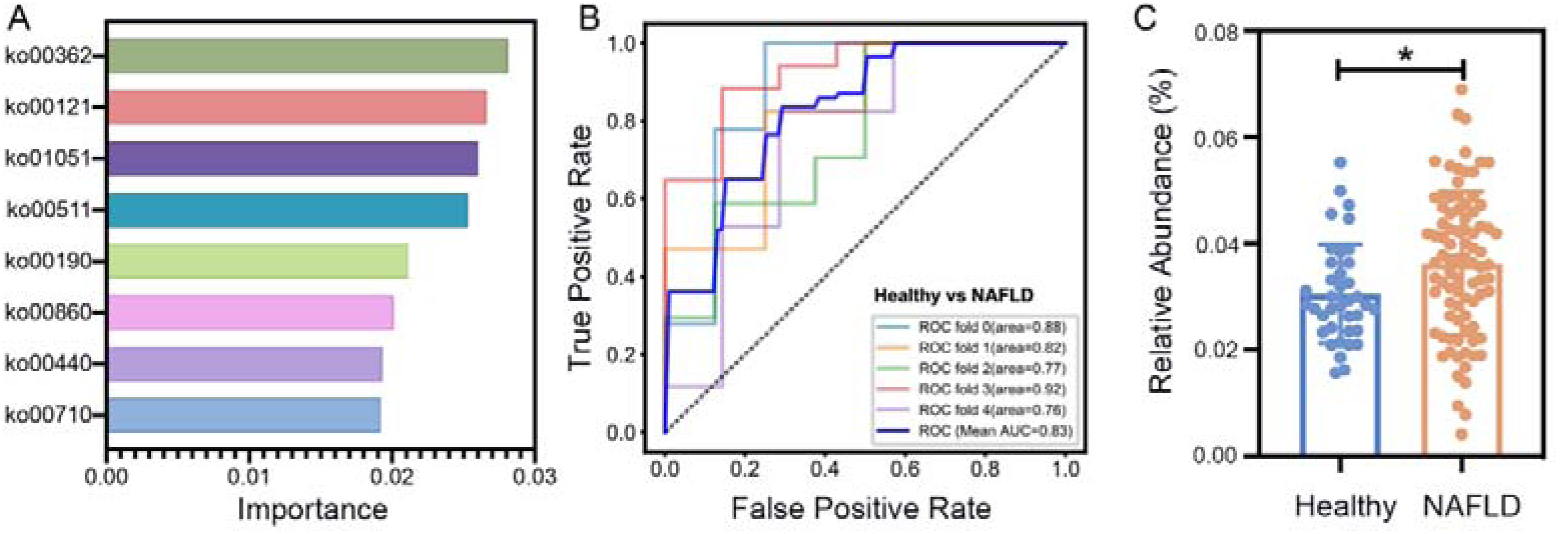
The differential pathway markers distinguishing NAFLD patients from healthy controls. Differential pathways were selected by two-tailed Mann-Whitney U- tests adjusted by Benjamini–Hochberg. Pathways with FDR values < 0.05 were included. Important differential pathway markers were then identified with random forest model and with the top 8 important pathways, the model achieved the highest AUC value. (A). The importance of pathways evaluated in NAFLD with the random forest model. (B). The AUC curve of random forest model with the top 8 important pathways. (C). The abundance of secondary A biosynthesis pathway (ko00121) in the healthy and the NAFLD groups. Values are the mean±SD. * indicates FDR<0.05.

### Novel genes and microbial genomes associated with secondary BA synthesis

The fact that the secondary BAs biosynthesis pathway was significantly elevated in NAFLD (Figure 3C) prompted us to examine the relevant BA metabolizing enzymes encoded by the microbiome. Taking advantage of the WMS data, we were able to quantify the gene abundance and to map these genes to specific microbial genomes.

#### Genes related to secondary BA synthesis

Bacterial genes directly involved in secondary BA synthesis catalyze the deconjugation, the oxidation and epimerization, or the multi-step 7α-dehydroxylation reactions (Figure 4A). Protein sequences of target enzymes were collected from Integrated Microbioal Genomes(IMG) database (Figure 4A).[18] High quality protein sequences were selected to construct hidden Markov models(HMMs), in order to identify potential BA metabolizing enzymes.

**Figure 4.**
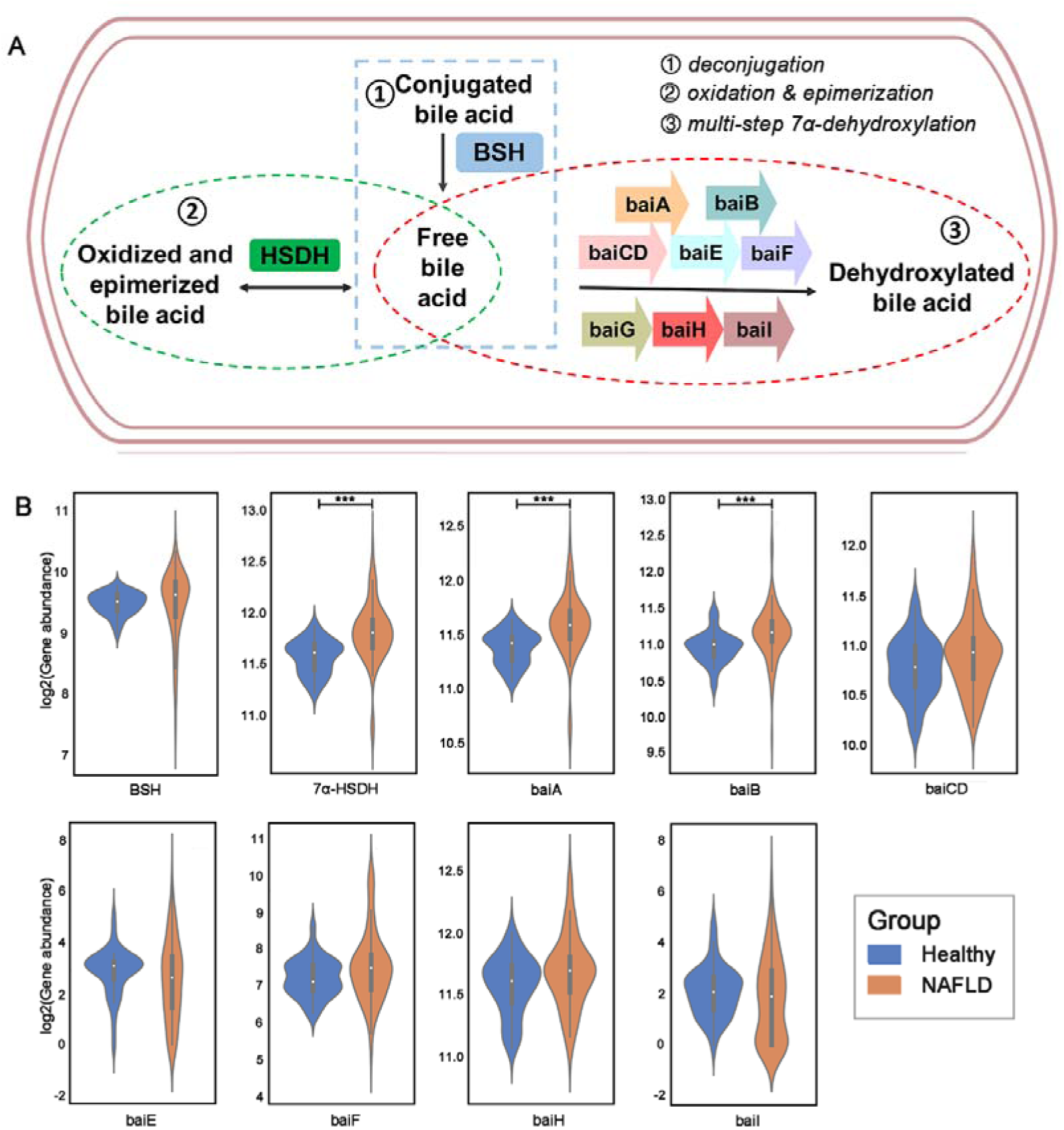
The abundance of the bacterial genes related to secondary bile acid synthesis. (A) Genes responsible for secondary bile acid biosynthesis can be grouped into 3 categories: (1) deconjugation, (2) oxidation and epimerization and multi-step 7α-dehydroxylation. (B) Gene abundance in health and NAFLD groups. Differences were identified by two-tailed Mann-Whitney U- tests adjusted by Benjamini–Hochberg. BSH: bile salt hydrolase; HSDH: hydroxysteroid dehydrogenase; baiA, 3α-hydroxysteroid dehydrogenase; baiB, bile acid-coenzyme A ligase; baiCD, 7α -hydroxy-3-oxo-D4-cholenoic acid oxidoreductase; baiE, bile acid 7α- dehydratase; baiF, bile acid coenzyme A transferase/hydrolase; baiG, primary bile acid transporter; baiH, 7beta-hydroxy-3-oxochol-24-oyl-CoA 4-desaturase; baiI, bile acid 7beta-dehydratase. *** indicates FDR<0.001.

The data (Figure 4B) showed that genes encoding 7-alpha-hydroxysteroid dehydrogenase(7α-HSDH), BSH and bile acid inducible operon (bai)A, baiB, baiCD, baiH were reletively more abundant than baiE, baiF and baiI. Importantly, significantly increased abundance of 7α-HSDH, baiA and baiB were observed in NAFLD compared to controls. These data were consistent with the pathway analysis results, and confirmed the increased secondary BA production in NAFLD.[12]

#### Novel identification of microbial genomes related to secondary BA synthesis using advanced bioinformatics

To identify the BA metabolizing microbial genomes, the metagenomic-assembled species(MAG) analysis was performed. Prevalent genes in the non-redundant gene catalog that presented in more than 5 samples were binned into 252 MAGs, which were considered to represent distinct microbial genomes. Among these, 50 MAGs that contain at least one gene encoding BSH, HSDH or bile acid inducible operons (Table S4) were defined as BA-metabolizing MAG. To obtain relatively complete microbial genomes, we re-assembled these 50 MAGs using high quality reads mapped to genes in each MAG.

Among these, 10 MAGs exhibited significantly increased abundance in NAFLD, while 18 MAGs were reduced in NAFLD (Figure 5A). Among the 10 MAGs elevated in NAFLD, 6 MAGs belong to Bacteroides (order Bacteroidales), including *B.vulgatus, B.ovatus*, and *B.stercoris*. Other MAG genomes were assigned as *E.rectale* and *E.biforme* (order Clostridiales). BA-metabolizing MAGs with reduced abundance in NAFLD are mainly from *R.bromii, D.longicatena* and *B. dorei*. Furthermore, we explored the species’ contributions of pathways in via HUMAnN2, and found that the pathway secondary bile acids biosynthesis were mainly encoded by *E.eligens* (48.3%) and *B.vulgatus* (26.2%) (Figure S5). This is consistent with the increased BA-metablizing MAGs belonging to species Bacteroides vulgatus and Eubacterium eligens.

**Figure 5.**
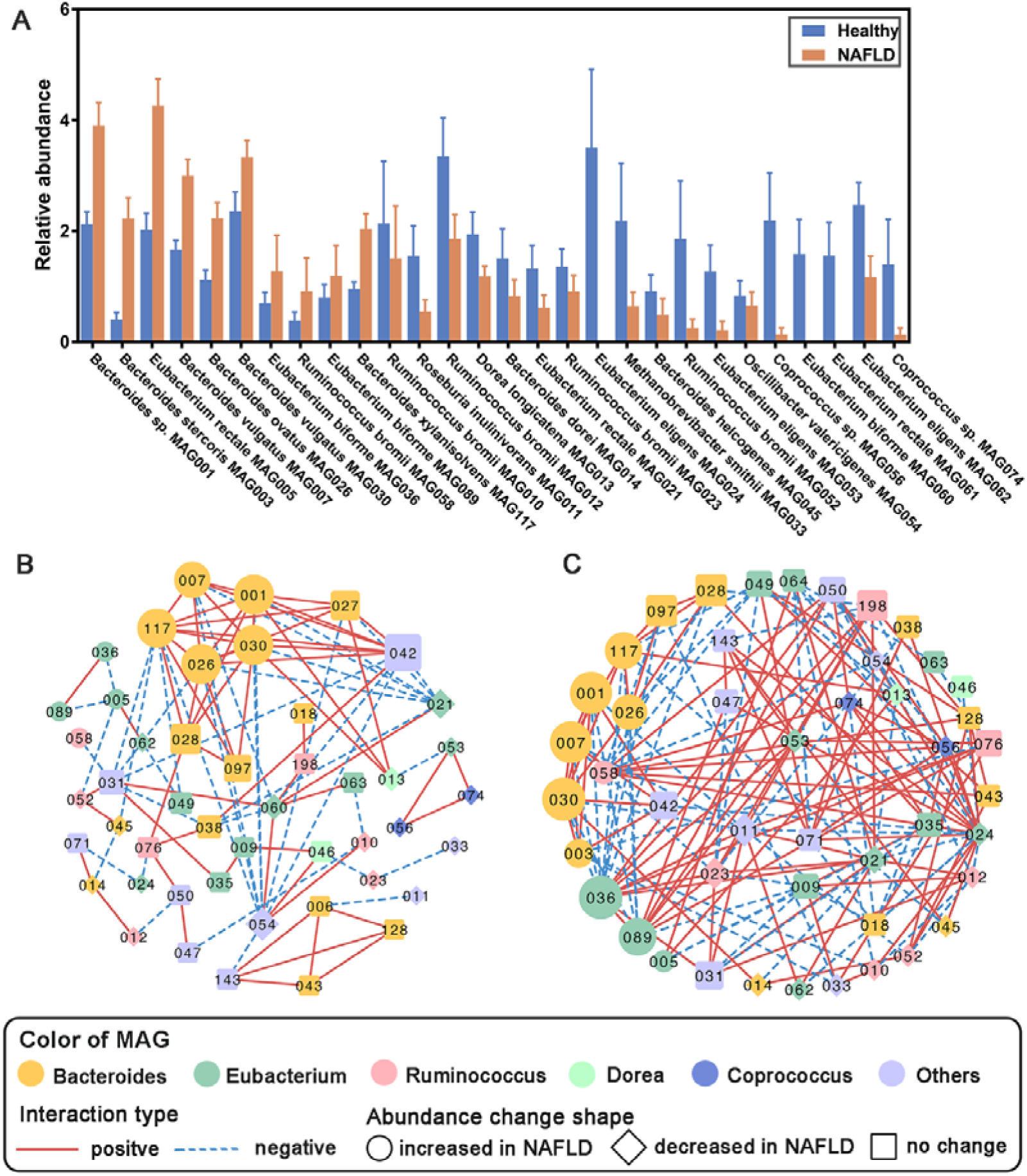
BA metabolizing MAG in NAFLD and healthy subjects. (A) MAG exhibiting differential abundance between healthy controls and NAFLD patients. Differential MAG were selected by two-tailed Mann-Whitney U- tests adjusted by Benjamini–Hochberg. MAG with FDR values < 0.1 were included. Values are mean ± SEM. Interaction network for BA metabolising MAG community in healthy controls (B) and NAFLD patients (C). Microbial interactions were calculated using SparCC with 100 refining interactions, and p value of each interaction is approximated with 1000 permutations. Only interactions with p value < 0.05 were included.

For a better understanding of the BA metabolizing microbial community, microbial interactions analysis was performed with BA-metabolizing MAGs. In contrast to the situation where more interactions existed in healthy group on whole-community level, we found that the sub-network of BA-metabolizing MAG was more complex with considerable interactions in NAFLD than in controls (164 and 100 edges, respectively) (Figure 5B &C). In addition, most MAGs with higher proportions in NAFLD patients were hub nodes in both healthy and NAFLD BA-metabolizing communities and were positively interacted, such as *Bacteroides sp*. MAG001, *B.vulgatus* MAG007, *B.ovatus* MAG026, *B.vulgatus* MAG030 and *B.xylanisolvens* MAG117. These are likely “house-keeping” species for BA metabolism. In contrast, Bacteroides stercoris MAG003, an MAG not included in the healthy network, was highly elevated in NAFLD, ranked high in the NAFLD network, and positively interacted with the “house-keeping” BA metabolizing species. Similarly, *E.biforme* MAG036 and MAG089, which exhibited the lowest hub score in healthy network, ranked the highest in NAFLD network.

In general, the observed species were represented by multiple MAGs. Here, *R.bromii* was represented by 7 MAGs, and *E.eligens* by 5 MAGs. However, only one of the 7 *R.bromii* MAG was significantly increased in NAFLD group, while 4 others showed decreased abundance (Table S5). Situations were similar in *B.vulgatus* (two of three increased) and *E.rectale* (one increased and two decreased). Unexpectedly, multiple MAGs of the same species were distributed in different modules both in healthy and NAFLD communities(Table S6). Apparently, these observations indicate that strains within the same species may function differently.

### Different BA metabolizing potentials among NAFLD microbiota and emergence of two subtypes of NAFLD: High BA versus normal BA subtype

Although the average abundances of the secondary BA metabolism pathway and related genes were increased in NAFLD, we noticed that the abundances exhibited a broad distribution among NAFLD patients (Figure 3C and 4B). Many of the NAFLD microbiota exhibited BA metabolizing potentials similar to those of healthy controls. Based on the abundance of 3 differential BA-metabolizing genes (7α-HSDH, baiA and baiB), NAFLD patients were clustered into two subtypes: normal-BA subtype comprising 45 patients and high-BA subtype comprising 37 patients (Figure 6A), which was not related to the disease severity (p=0.7). The abundances of the 3 marker genes were all significantly higher in high-BA subtype, but were similarly represented between normal-BA subtype and healthy control group (Figure 6B). In addition, we performed the PCA analysis based on the entire differential microbial enzymes and found that the normal-BA subtype and the healthy control group exhibited closer distance, as compared to the high-BA group (Figure 6C). In further characterization of the microbial profiles of the patterns of the normal-BA and high-BA groups, we identified 3 species (Table S7), 68 enzymes (Table S8) and 16 pathways (Table S9) that could distinguish the normal-BA subtype from the high-BA subtype, and, at the same time, could distinguish NAFLD from the healthy group. Based on the relative abundance of these differential features, the study subjects were clustered into three groups consistent with their BA metabolizing potentials. Features were also clustered into two groups (Figure S6). One group (including species Flavonifractor plautii, enzymes 2-dehydropantoate 2-reductase and glutamate 5-kinase and pathway glycosaminoglycan degradation etc.) exhibited elevated abundance in normal-BA subtype and reduced abundance in high-BA subtype. The other group (including species Escherichia coli and Ruminococcus bromii, enzymes glycerol dehydrogenase, agmatinase and pathway citrate cycle, phosphotransferase system etc.) exhibited an opposite distribution among the study groups.

**Figure 6.**
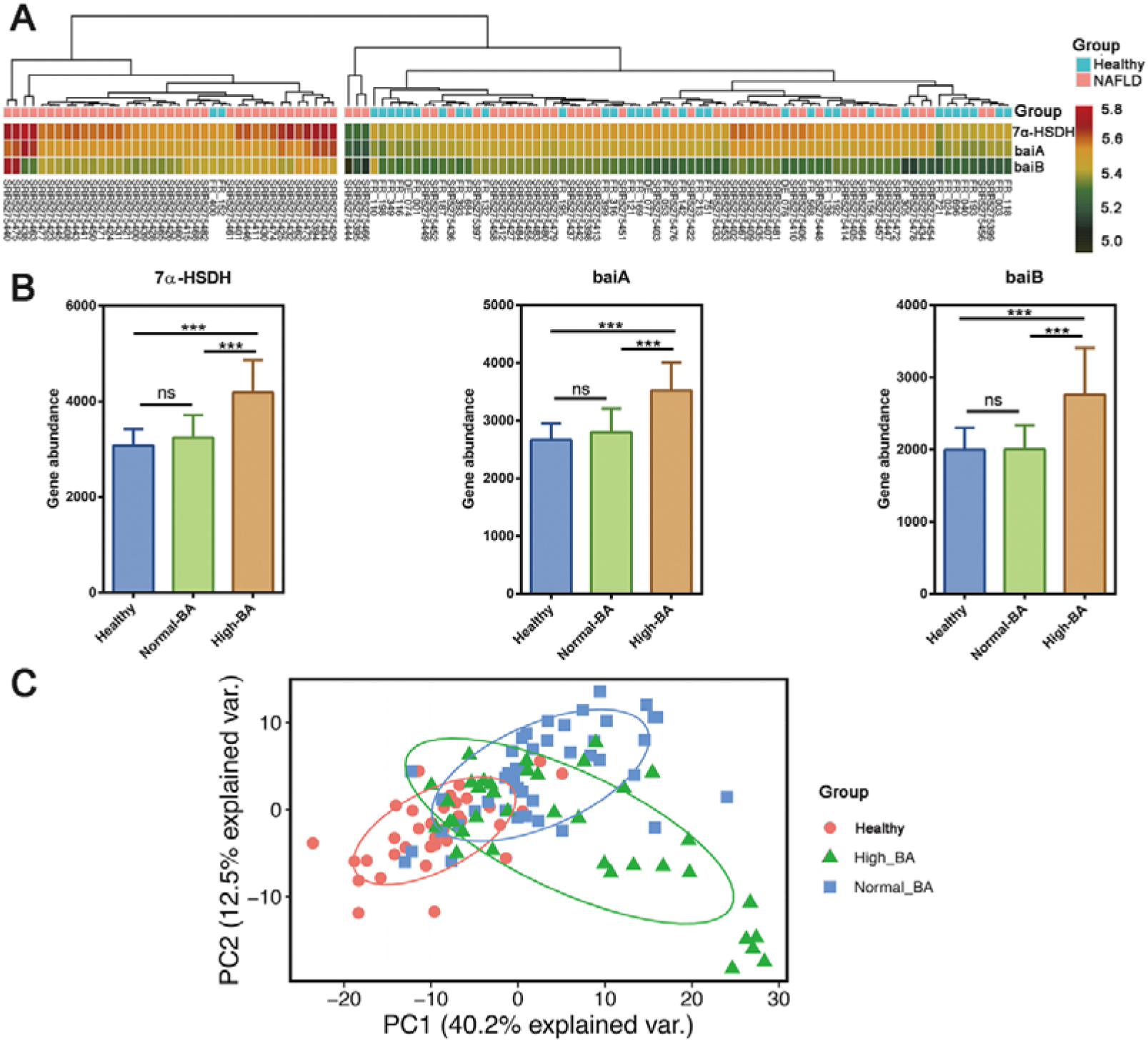
Subgroups of NAFLD patients with different abundances of the secondary BA synthesis genes. (A) NAFLD patients were clustered into two subgroups: normal-BA subgroup and high-BA subgroup according to the abundances of 3 differential secondary BA synthesis genes. (B) Comparison of the abundances of 3 differential secondary BA synthesis genes among healthy control, normal-BA and high BA groups. They were all significantly increased in high-BA subgroup, but was not different between normal-BA subgroup and healthy group (Dunn tests adjusted by Benjamini–Hochberg). (C) PCA plot based on the differential enzymes. Subjects were clustered according to the secondary BA metabolizing potentials (p <0.001 with ANOSIM analysis). Values are mean±SD. *** indicates FDR<0.001.

### Elevated secondary BA synthesis capability in the validation cohort of NAFLD

Similar analyses were performed with the validation dataset. The secondary BA synthesis genes 7α-HSDH, BSH,baiA, baiB, baiCD, baiF, and baiH were reletively more abundant than baiE and baiI. Importantly, significantly increased abundance of most secondary BA synthesis genes were observed in NAFLD compared to controls (Figure S7).

As for BA metabolizing microbial genomes, we identified 13 MAGs, each carrying at least one gene encoding BSH, HSDH or bai operon. Among these, 9 MAGs exhibited a trend of increased abundance in NAFLD. Consistent with the discovery cohort, these 9 MAGs belonged to *B.vulgatus*, and *R. bromii*(Table S10). Statistical significance was not achieved for the increased abundances of the MAGs, likely due to the small sample size.

### Discussion

In this study, we defined the structural and functional differences in gut microbiota between NAFLD and healthy subjects, at the resolutions of gene, species and strain. The current study is novel in using WGS data to compare the gut microbiota between NAFLD and healthy controls and underpinning the role of BA metabolizing microbiome in NAFLD, and potentially identifying two microbiota-derived subtypes of NAFLD that may have clinical implications for both biomarker as well as therapeutic development. Compared with the approach of 16S rRNA sequencing, WMS data allow direct function quantification and accurate taxa assignment of the entire gut microbiome, at the levels of species and strain. Out of the many differential representations of genes and species between NAFLD and healthy controls, one outstanding observation is the increased abundance of secondary BA metabolizing genes and microbes in NAFLD and that BA metabolizing bacteria were dominant taxa in the gut of NAFLD. For the first time, we identified the genes and bacterial strains responsible for elevated secondary BA synthesis in NAFLD. Similarly, increased abundances of the BA metabolizing genes and bacterial species were observed in an independent validation cohort. Considering the profound impact of BA signaling on lipid and carbohydrate metabolism[19], the differential BA metabolizing genes and bacterial strains we identified may serve as novel therapeutic targets for NAFLD management.

We and others have reported elevated secondary BA production in NAFLD. [12, 20] In our previous study[12], we observed much increased secondary BAs in NAFLD serum and consistently, an elevated taurine metabolizing microbiota, an indication of increased BA metabolism in the gut. However, we did not observe any significant change in the abundance of those microbes that directly metabolize BA (that is, microbes encoding BSH, 7-alpha-HSDH and 7-alpha-dehydroxylase), likely because the 16S rRNA sequencing approach was not able to provide a sufficient resolution for functional analysis. With the advantage WGS data, the current study was able to provide convincing evidence at a satisfactory resolution, that secondary BA synthesis enzymes and microbes with secondary BA metabolizing potentials were indeed elevated in NAFLD gut microbiota. As secondary BAs are potent antagonistic ligands for FXR, data presented here is a strong support for the hypothesis that elevated secondary BA synthesis by the microbiota contributes to NAFLD etiology.[12, 21]

Although on average NAFLD patients exhibited elevated BA metabolizing microbiota, and higher serum DCA (secondary BA) when compared to healthy controls, our data showed that elevated BA metabolizing microbiota was not a unanimous phenomenon in NAFLD. More than half of the NAFLD patients (45 out of 82) had a microbiota with normal BA metabolizing potential. Based on BA metabolizing potentials, our NAFLD patients can be clustered into two subtypes. This indicates that BA related pathomechanism does not apply to many NAFLD patients, in line with the current multi-hit hypothesis.[3] Besides the difference in BA metabolizing potentials, these two subtypes of the gut microbiota also exhibit different abundances in other genes, pathways, and bacterial species. It is interesting to note that NAFLD microbiota with higher BA metabolizing potentials also exhibited elevated representation of *E.coli*, a potent alcohol producer[6, 22], suggesting that the gut microbiota may impact NAFLD pathogenesis through multiple mechanisms in the same patient.

BA based therapies such as obeticholic acid has been shown to improve NASH. [23] However, the response rates to OCA in improvement of one-stage of fibrosis in the FLINT trial was 35% versus 19% in placebo.[24] It is plausible that NAFLD patients with altered BA subtype may be more likely to respond to BA based therapies and those with a normal BA subtype should receive an alternate strategy paving the pay for a microbiome based precision medicine tool in NASH therapeutics.

Another outstanding observation in this study is that many strains of the same species are functionally different. Specifically, different strains of Bacteroides ovatus were clustered into different functional modules (modules 0, 2, 4 in healthy communities and modules 3, 4, 6 in NAFLD communities). It is also interesting to note that only one of the four observed strains of Bacteroides ovatus was significantly increased in NAFLD group. Similar observations were reported for *F. prausnitzii*[25, 26] and *E.coli*[27, 28], suggesting the genomic variability within a microbial species.[29] Some of the microbiome studies based on 16S rRNA platforms may need a re-evaluation because of this genomic variability.

It was interesting to note that 10 BA-metabolizing bacterial strains, including *B.stercoris, E.biforme*, and *R.bromii*, were elevated and were dominant strains in NAFLD microbiota. These BA-metabolizing strains belong to two different phylum. Zhao et al. proposed a concept in gut microbiota that a group of species that “exploit the same class of environmental resources in a similar way” may be considered as a “guild” in ecology[30] and members of a guild do not necessarily share taxonomic similarity, but they co-occur when adapting to the changing environment.[25] Similarly, the 10 BA-metabolizing strains may act as a synergetic guild to promote the secondary BA production in the NAFLD microbial community. There were more positive interactions among these 10 strains in NAFLD community than in healthy community, indicating elevated capabilities of secondary BA production and intensified competition among these secondary BA producers within the microbial guild of NAFLD. It is likely that these strains are responsible for elevated secondary BA production in NAFLD, contributing to NAFLD pathogenesis.[12] Among these 10 strains, MAG036,MAG089,and MAG003 with increased abundance and the highest network importance in NAFLD may act as the “keystone” species[53], and therefore, represent potential targets for intervention.

At the whole community level, the NAFLD gut microbiota exhibited significantly reduced diversity compared to the healthy controls. In addition, much reduced interactions among the members of the NAFLD gut microbiota were observed. With less strains and sparse interactions, the gut microbial community in NAFLD is relatively weak and unstable. Similarly, reduced biodiversity were reported in the gut of obesity.[31] It is postulated that long-term dietary habit is the major cause for the altered gut microbiota.[32] The biodiversity disaster in the gut of humans demands immediate attention. The restoration of the gut microbial diversity may, at the same time, prevent or cure many of the microbiota related diseases including NAFLD.

In summary, we identified specific genes and bacterial strains responsible for elevated secondary BA production in NAFLD. These genes and strains may serve as novel therapeutic targets for microbiome-based high-BA subtype of NAFLD. These findings strongly support our hypothesis that elevated secondary BA synthesis contributes to the development of NAFLD. In addition, our WGS study revealed the heterogeneity of the gut microbiota among NAFLD patients highlighting the importance of personalized treatment for NAFLD. Our study also revealed many other microbial characteristics of the NAFLD that demands attention such as the much reduced diversity and the ecological guild in the gut of NAFLD.

## Materials and Methods

### Data information and preprocessing

Disco very dataset: The NAFLD datasets and relevant meta data(Sequence Read Archive, PRJNA373901) were described previously[9] comprising 86 biopsy-proven NAFLD patients. The healthy control dataset was from PRJEB6070[33], with 38 healthy individuals with BMI < 25. These subjects were chosen because of similar age and gender ratio compared to NAFLD patients to effectively reduce bias[34] (Table 1 & Table S1).

Validation dataset: 10 middle-aged NAFLD subjects [35] (PRJNA420817) were recruited to a diet trial and the initial baseline data before diet interventionwere used for this study. 11 healthy subjects from MetaHit Project[36](Sequence Read Archive, PRJEB1220) with similar age and gender ratio were chosen as controls (Table 1& Table S1).

All subjects provided a written informed consent and the study protocol was approved by Institutional Review Board (approval number:UCSD IRB11298) or registered at ClinicalTrials.gov with identifier: NCT02558530.

The KneadData(http://huttenhower.sph.harvard.edu/kneaddata) tool was used to ensure the data consisted of high quality microbial reads free from contaminants. The low quality reads were removed using Trimmomatic(SLIDINGWINDOW:4:15 MINLEN:75 LEADING:10 TRAILING:10). The remaining reads were mapped to the human genome(hg38) by bowtie2[37], and the matching reads were removed as contaminant reads from the host.

### Gene-based taxonomic and functional profiling of gut microbiota

MetaPhlAn2[38] was used to identify the composition of gut microbial community and to assess the abundance of the prokaryotes within each sample. Species that failed to exceed 0.01% relative abundance in at least 20% samples were excluded.

The functional profiling of gut microbiome was determined by the HMP Unifiled Metabolic Analysis Network (HUMAnN2)[39]. In brief, high-quality metagenomic reads were mapped to the pangenomes of species identified with MetaPhlAn2 and these pangenomes have been pre-annotated by UniRef90 families. Reads failed to map to a pangenome were aligned to UniRef90 by translated search with DIAMOND[40]. Hits to UniRef90 are weighted according to alignment quality, sequence length and coverage. In this study, enzyme abundance was quantified by regrouping (summed) according to EC number and pathway abundance by regrouping (summed) genes in pathways against KEGG database.

### Identification of genes required for secondary BA synthesis

To identify genes that encode enzymes catalyzing secondary BA synthesis, hidden Markov models (HMMs) of BA-related genes were constructed. Secondary BA synthesis mainly involves (1) deconjugation, (2) oxidation and epimerization and (3) multi-step 7α-dehydroxylation. Enzymes participating in these processes are bile salt hydrolase (BSH), hydroxysteroid dehydrogenase (HSDH) and enzymes required in the multi-step 7α-dehydroxylation (including baiA, baiB, baiCD, baiE, baiF, baiH and baiI).[18] Representative protein sequences of target enzymes were obtained from Integrated Microbioal Genomes (IMG) database[41]. High quality sequences were selected and aligned in Clustal Omega[42] before they were used to construct HMMs on full-length proteins via hmmbuild in HMMER(3.1b2)[43]. Model seed sequences were realigned to the model using hmmalign (default mode) before rebuilding models based on the obtained alignments until both model length and relative entropy per position were constant. Subsequently, all protein sequences in non-redundant gene catalog were screened (hmmsearch) for candidate protein sequences and sequences with hmmscore > lower quartile score and e-value less than 10-5 were identified as potential secondary BA synthesis associated genes.

### Assembly-based microbial genomes

For functional analysis of the microbial genomes, we performed bin-based microbial genome assembly with the WMS data, including de nove assembly and non-redundant human gut gene catalog construction, co-abundance clustering and determination of metagenome-assembled genomes (MAG), MAG-augmented assembly and taxonomic annotation.

#### De novo assembly and non-redundant human gut gene catalog construction

High-quality paired-end reads from each sample were used for de novo assembly with Megahit[44] into contigs of at least 500-bp length. Genes were predicted on the contigs with MetaGeneMark[45]. A non-redundant gene catalog related to NAFLD was constructed with CD-HIT[46] using a sequence indentity cut-off of 0.95, with a minimum coverage cut-off of 0.9 for the shorter sequences and 11,348,567 microbial genes were contained.

#### Co-abundance clustering and determination of MAG

Bowtie2 was used to align high quality reads to the non-redundant gene catalog. Aligned results were random sampled and downsized to 15 million per sample (FR-173, FR-719, FR-730, SRR4275396, SRR4275459, SRR4275469, SRR4275470 were excluded for not enough reads) to adjust for sequencing depth and technical variability. The soap.coverage script (available at: http://soap.genomics.org.cn/down/soap.coverage.tar.gz) was used to calculate gene-length normalized base counts and the gene abundance profiling was calculated as the average abundance of 30 times of repeated sampling. All the genes were clustered into MAG using MSPminer[47] based on their abundance with default parameters.

#### MAG-augmented assembly and taxonomic annotation

We performed augmented assembly for target MAG. Briefly, the MAG- and sample-specific reads were derived by aligning all high-quality reads to the MAG gene contigs with Burrows-Wheeler Aligner (0.7.17)[48], followed by de novo assembly with SPAdes(3.13.0)[49] using k-mers from 21 to 55. CVtree3.0 web server[50] was used to identify the taxonomy of the MAGs, which applies a composition vector to perform phylogenetic analysis.

### Statistic analysis

#### Differential features identification

Compositional features and functional features that present in at least 20% of the samples and with average relative abundance over 0.01% in each group were selected for further differential analysis. Differential features were identified by two-tailed Mann-Whitney U-tests adjusted by Benjamini-Hochberg. Features with an FDR value < 0.05 (FDR values < 0.1 for species) were identified as differential features. Then differential compositional and functional feature profiles were used to build random forest(RF) model using RandomForest package in R. Feature importance were estimated via gini importance and then the best model were rebuilt by adding features according to their importance ranks. Area Under the Receiver-Operator Curve(AUC) was used to measure the accuracy of the models.

#### Microbial interaction analysis

SparCC[51] was performed to construct compositionality-corrected microbial interactions network, which is capable of estimating correlation values from compositional data. Interactions were calculated with 100 refining interactions, after which statistical significance of each interaction was estimated with 1000 permutations. Only interactions with p value < 0.05 were included in downstream analysis and those interactions with magnitudes > 0.4 were included in the “core community”. The importance of species in the community was calculated using Hyperlink-Induced Topic Search(HITS) algorithms in Python package ‘networkx’. The networks were then visualized with Cytoscape[52] and module analysis was performed with ModuLand in Cytoscape.

#### Other statistics

Analysis of similarities (ANOSIM) was performed based on distance matrix for statistical comparisons of samples between groups or subtypes. P value was calculated using 9999 permutations. p < 0.05 indicates significant difference. Hetamap was plotted via “pheatmap” package in R, and features were clustered based on euclidean distance by “ward.D”. Differential features among healthy, normal-BA and high-BA groups were identified with Dunn tests adjusted by Benjamini–Hochberg, and features with FDR values < 0.05 were determined as significant differential features.

## Abbreviations

baiA: 3α-hydroxysteroid dehydrogenase;
baiB: bile acid-coenzyme A ligase;
baiCD: 7α
-hydroxy-3-oxo-D4-cholenoic acid oxidoreductase;
baiE: bile acid 7α- dehydratase;
baiF: bile acid coenzyme A transferase/hydrolase;
baiG: primary bile acid transporter;
baiH: 7beta-hydroxy-3-oxochol-24-oyl-CoA 4-desaturase;
baiI: bile acid 7beta-dehydratase;
BAs: bile acids;
BSH: bile salt hydrolase;
FXR: farnesoid X receptor;
HMM: hidden Markov model;
HSDH: hydroxysteroid dehydrogenase;
MAG: metagenome-assembled genome;
NAFLD: non- alcoholic fatty liver disease;
NASH: non-alcoholic steatohepatitis;
WMS: whole metagenome sequences.

## Disclosures

The authors have declared that no competing interests exist.

## Availability of data and materials

The datasets supporting the conclusions of this article are available in the NCBI’s Sequence Read Archive repository (https://www.ncbi.nlm.nih.gov/bioproject/), under study accession number PRJNA373901, PRJNA420817, PRJEB1220 and PRJEB6070.

## Synopsis

The microbial markers identified at the species/strain levels may be useful for non-invasive diagnosis of NAFLD. The microbial differences in bile acid metabolism and strain-specific differences among NAFLD microbiota highlight the potential for precision medicine in NAFLD treatment.

## Notes

Grant support: This work was supported by National Natural Science Foundation of China 81774152 (to RZ), 81770571 (to LZ), National Postdoctoral Program for Innovative Talents of China BX20190393 (to NJ), China Postdoctoral Science Foundation 2019M663252 (to NJ) and 2019M651568 (to DW), Natural Science Foundation of Shanghai 16ZR1449800 (to RZ), Fundamental Research Funds for the Central Universities 19ykzd01(to LZ) and 20kypy07(to NJ), the Guangzhou Science and Technology Plan Projects 201803040019 (to PL), Guangdong Province “Pearl River Talent Plan” Innovation and Entrepreneurship Team Project (2019ZT08Y464 to LZ) and the National Key Clinical Discipline of China, and Funds from the University at Buffalo Community of Excellence in Genome, Environment and Microbiome (GEM) (to LZ). RL receives funding support from NIEHS (5P42ES010337), NCATS (5UL1TR001442), and NIDDK (R01DK106419). The funders had no role in study design, data collection and analysis, decision to publish, or preparation of the manuscript.

### Competing Interest Statement

The authors have declared no competing interest.

