## Supplementary materials for "Alterations in bile acid metabolizing gut microbiota and specific bile acid genes as a precision medicine to subclassify NAFLD"

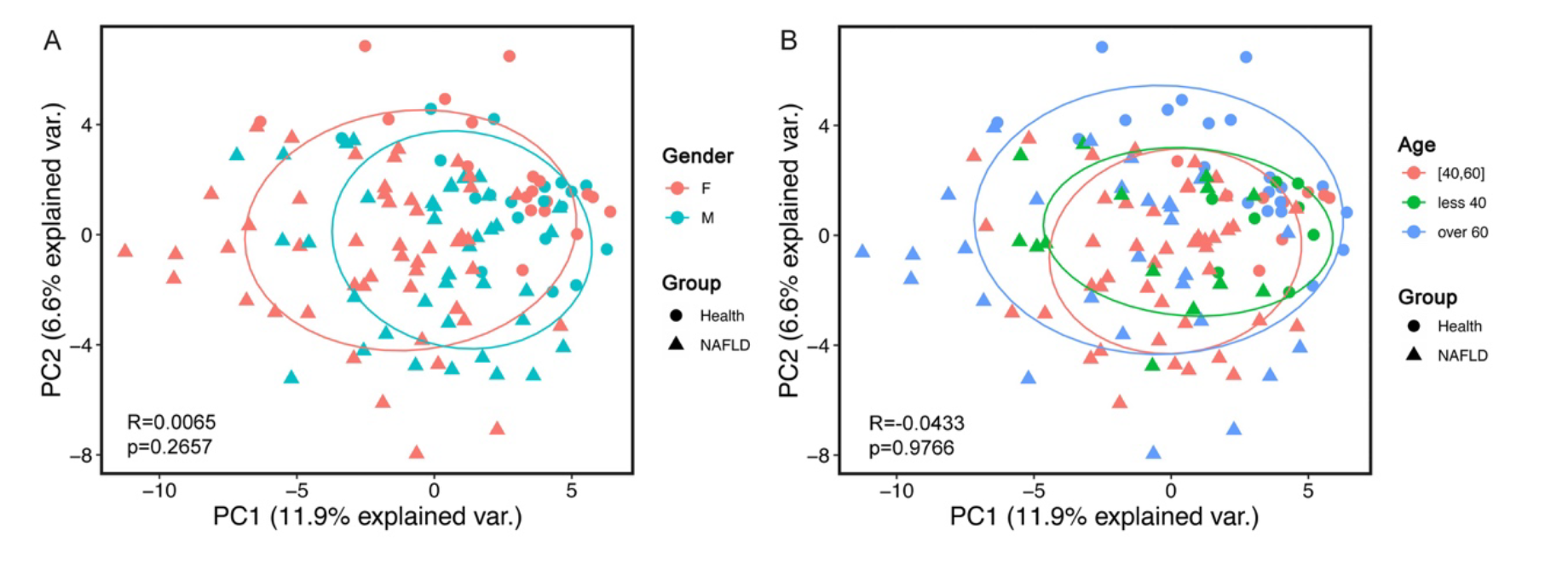
Figure S1 Principle component analysis based on Bray-Curtis dissimilarity of gender and age

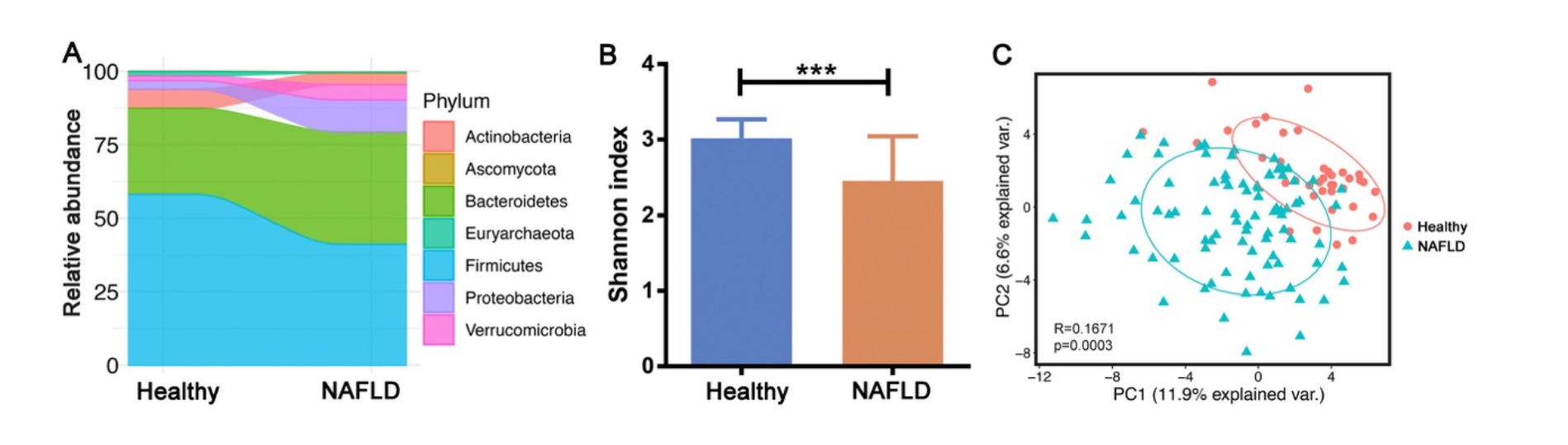
Figure S2 The gut microbiota composition of NAFLD patients and healthy controls

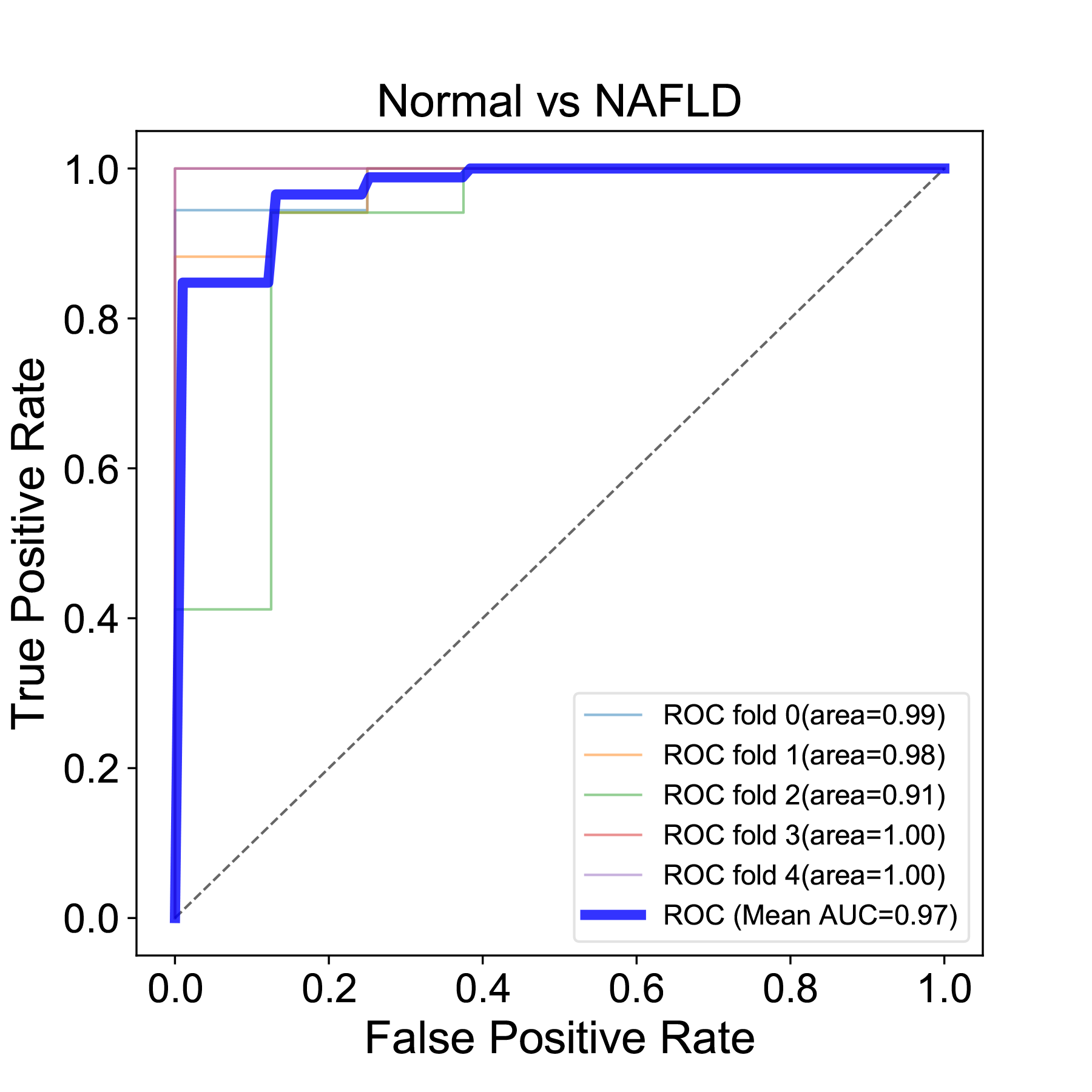

Figure S3 The AUC curve of random forest model with the differential species

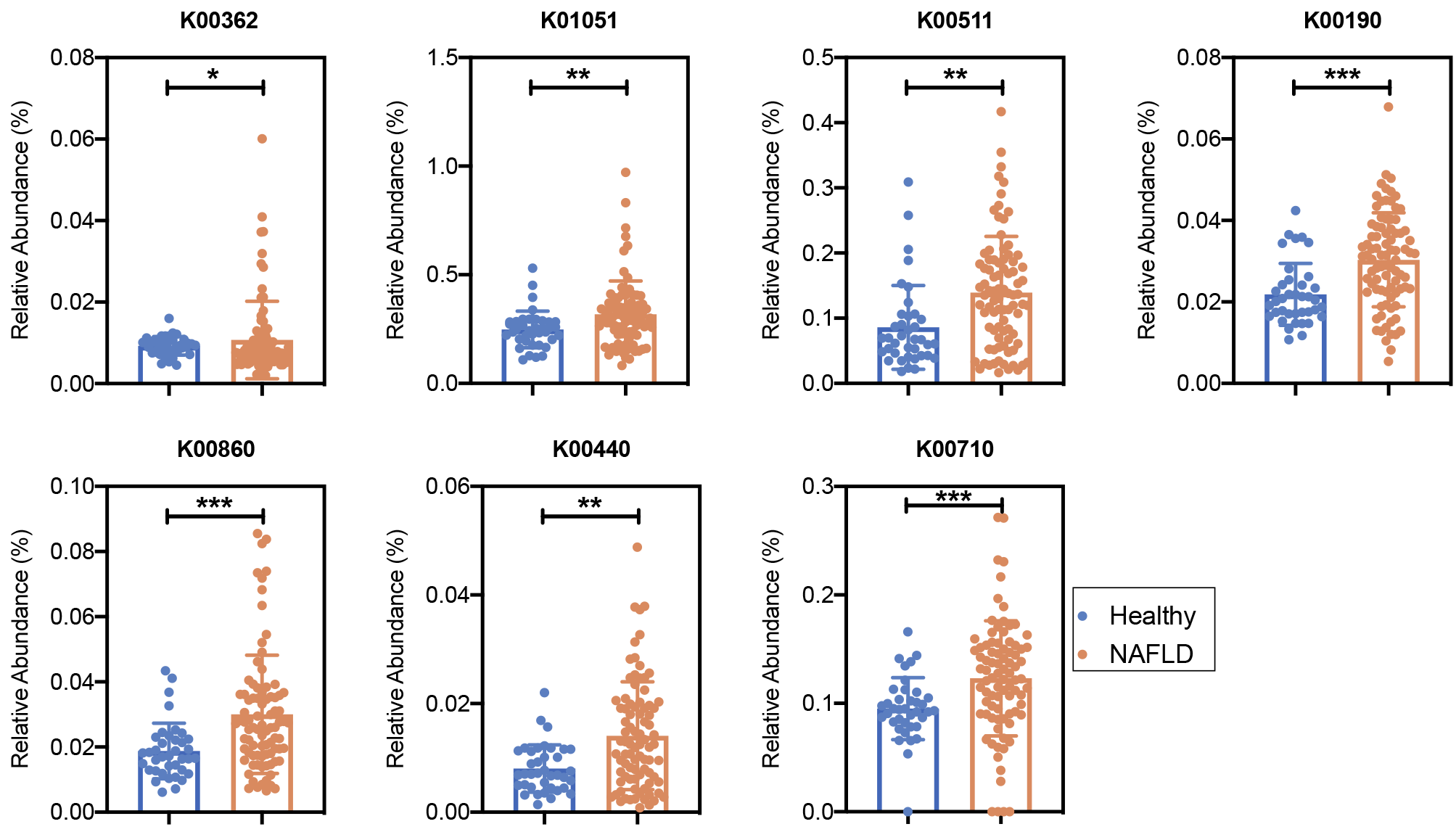
Figure S4 The relative abundance of important pathway features. ko00362: Benzoate degradation; ko01051: Biosynthesis of ansamycins; ko00511: Other glycan degradation; ko00190: Oxidative phosphorylation; ko00860: Porphyrin and chlorophyll metabolism; ko00440: Phosphonate and phosphinate metabolism; ko00710: Carbon fixation in photosynthetic organisms

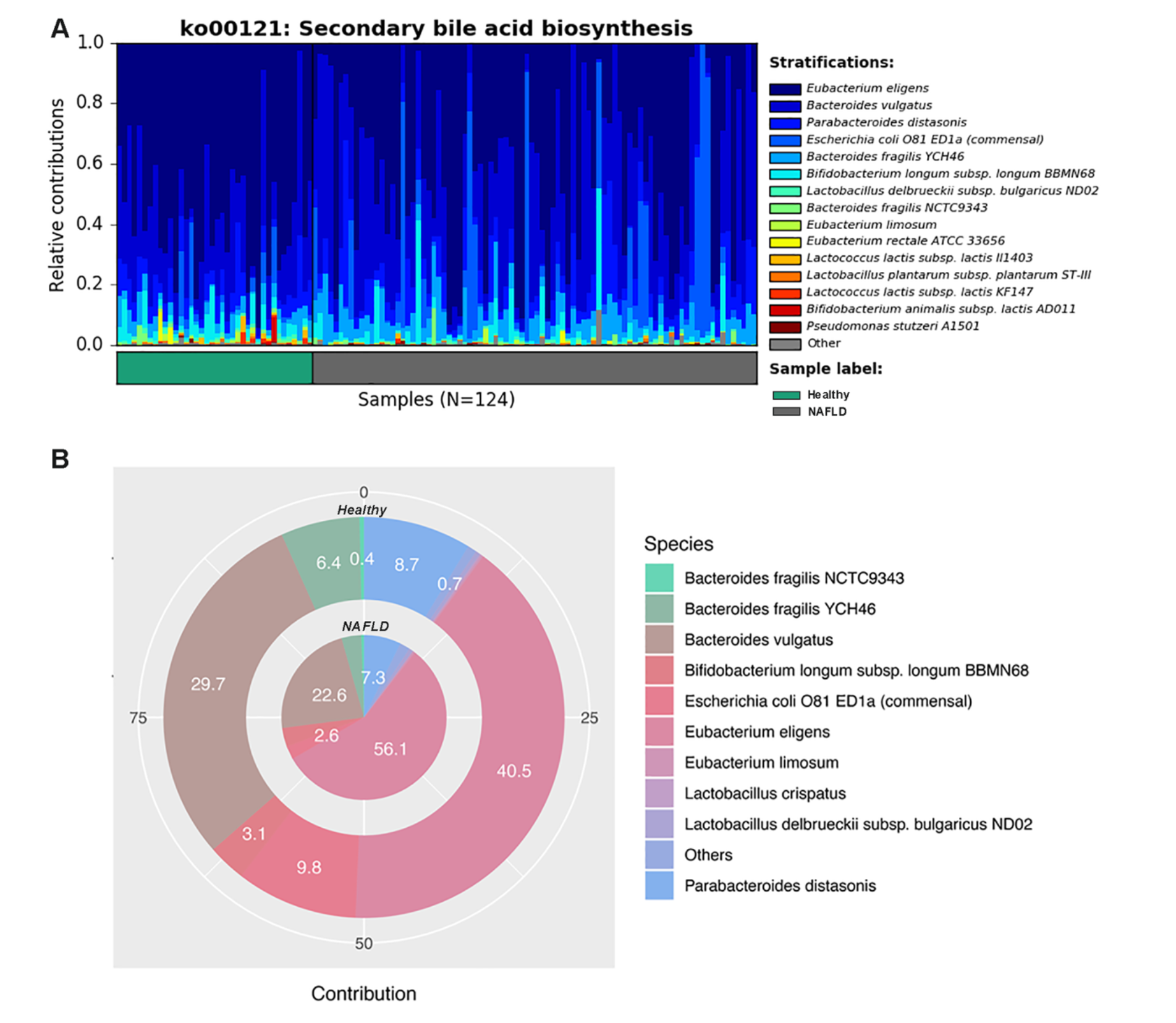
Figure S5 Species’ metagenomic contributions to secondary bile acid biosynthesis (ko00121)

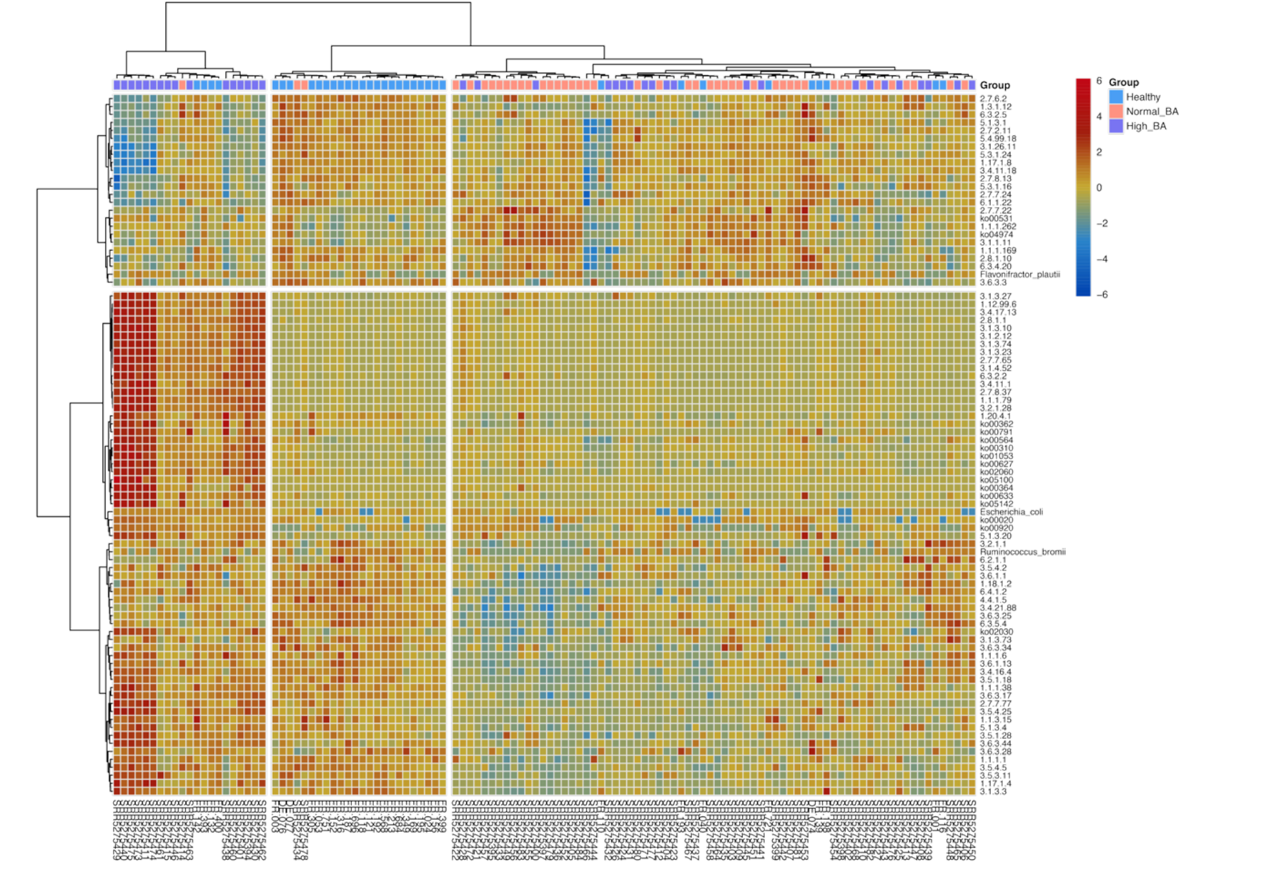
Figure S6 Microbiota diversities among NAFLD patients

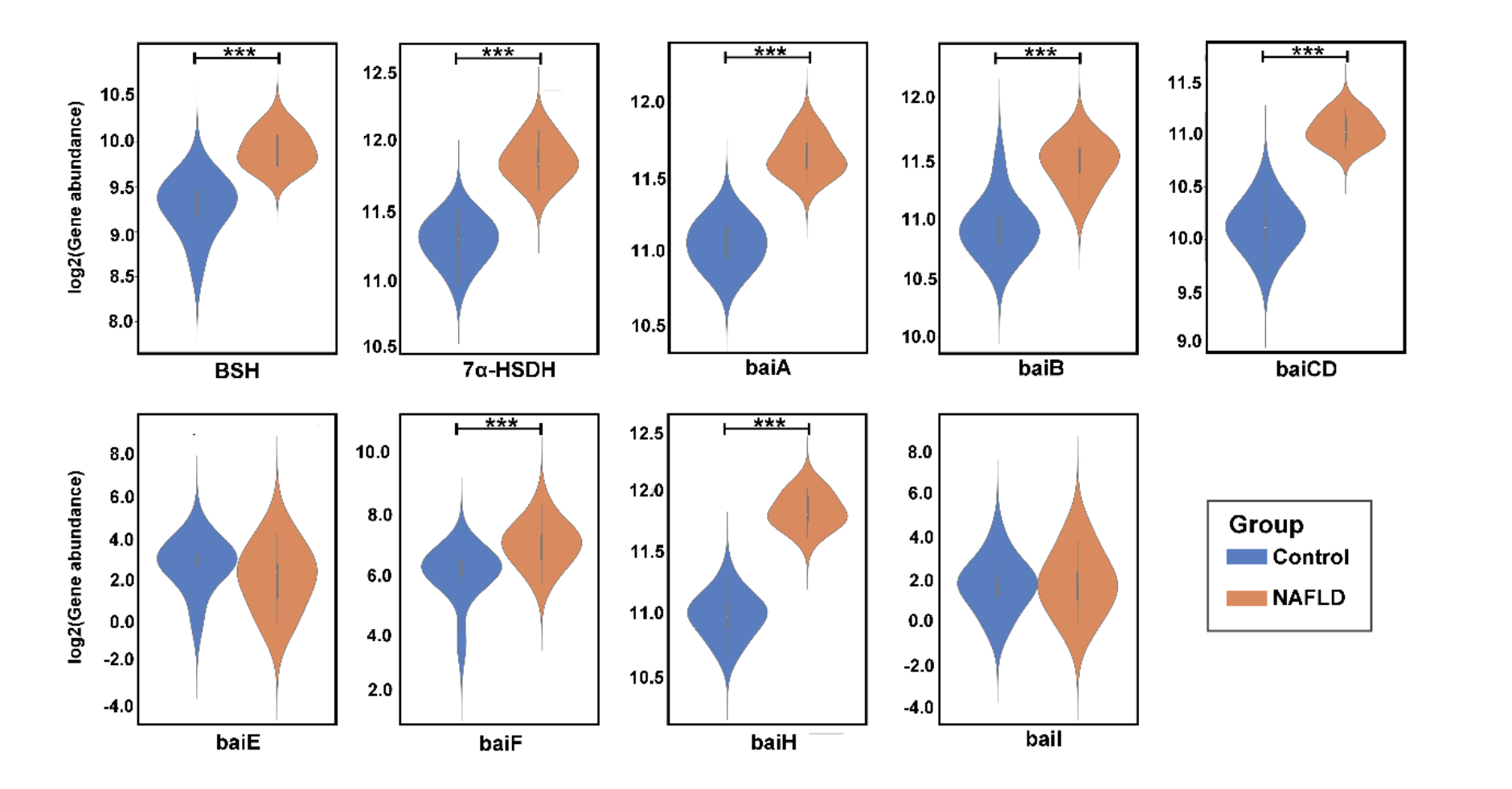
Figure S7 The abundance of the bacterial genes related to secondary bile acid synthesis in the validation cohort

| **Table S1 Dataset information and metadata** | | | | | | | |
| --- | --- | --- | --- | --- | --- | --- | --- |
| **Sample_name** | **Gender** | **BMI** | **Age** | **Status** | **Fibrosis stage** | **Country** | **Corhort** |
| SRR5275394 | F | 41.1 | 58 | NAFLD | 1 | USA | Discovery |
| SRR5275395 | M | 28.3 | 47 | NAFLD | 0 | USA | Discovery |
| SRR5275396 | F | 21.3 | 53 | NAFLD | 2 | USA | Discovery |
| SRR5275397 | M | 28.2 | 51 | NAFLD | 0 | USA | Discovery |
| SRR5275398 | F | 34.3 | 53 | NAFLD | 0 | USA | Discovery |
| SRR5275399 | M | 33.5 | 43 | NAFLD | 1 | USA | Discovery |
| SRR5275400 | M | 30.7 | 22 | NAFLD | 1 | USA | Discovery |
| SRR5275401 | F | 37.4 | 29 | NAFLD | 0 | USA | Discovery |
| SRR5275402 | M | 40.2 | 57 | NAFLD | 1 | USA | Discovery |
| SRR5275403 | M | 24.3 | 36 | NAFLD | 0 | USA | Discovery |
| SRR5275404 | F | 26.5 | 43 | NAFLD | 0 | USA | Discovery |
| SRR5275405 | F | 27.8 | 54 | NAFLD | 0 | USA | Discovery |
| SRR5275406 | M | 35 | 45 | NAFLD | 1 | USA | Discovery |
| SRR5275407 | M | 28.8 | 62 | NAFLD | 0 | USA | Discovery |
| SRR5275408 | M | 32.9 | 58 | NAFLD | 1 | USA | Discovery |
| SRR5275409 | M | 28.1 | 67 | NAFLD | 1 | USA | Discovery |
| SRR5275410 | M | 32.2 | 51 | NAFLD | 1 | USA | Discovery |
| SRR5275411 | M | 30.1 | 65 | NAFLD | 0 | USA | Discovery |
| SRR5275412 | M | 26.8 | 44 | NAFLD | 0 | USA | Discovery |
| SRR5275413 | F | 31.4 | 34 | NAFLD | 1 | USA | Discovery |
| SRR5275414 | F | 35.2 | 64 | NAFLD | 4 | USA | Discovery |
| SRR5275415 | F | 41.7 | 44 | NAFLD | 0 | USA | Discovery |
| SRR5275421 | F | 25.7 | 60 | NAFLD | 4 | USA | Discovery |
| SRR5275422 | F | 30.1 | 64 | NAFLD | 2 | USA | Discovery |
| SRR5275423 | F | 27.9 | 47 | NAFLD | 0 | USA | Discovery |
| SRR5275424 | M | 24.5 | 65 | NAFLD | 1 | USA | Discovery |
| SRR5275425 | F | 37.2 | 56 | NAFLD | 1 | USA | Discovery |
| SRR5275426 | M | 38.6 | 43 | NAFLD | 2 | USA | Discovery |
| SRR5275427 | M | 33.1 | 65 | NAFLD | 2 | USA | Discovery |
| SRR5275428 | F | 27 | 63 | NAFLD | 1 | USA | Discovery |
| SRR5275429 | F | 24.9 | 52 | NAFLD | 1 | USA | Discovery |
| SRR5275430 | F | 27.5 | 49 | NAFLD | 1 | USA | Discovery |
| SRR5275431 | F | 28.3 | 54 | NAFLD | 0 | USA | Discovery |
| SRR5275432 | F | 29 | 61 | NAFLD | 3 | USA | Discovery |
| SRR5275433 | M | 29.8 | 29 | NAFLD | 0 | USA | Discovery |
| SRR5275434 | F | 25.5 | 59 | NAFLD | 1 | USA | Discovery |
| SRR5275435 | M | 26.7 | 48 | NAFLD | 0 | USA | Discovery |
| SRR5275436 | F | 38.4 | 60 | NAFLD | 3 | USA | Discovery |
| SRR5275437 | M | 33.2 | 58 | NAFLD | 1 | USA | Discovery |
| SRR5275438 | F | 38.8 | 65 | NAFLD | 4 | USA | Discovery |
| SRR5275439 | F | 35.4 | 31 | NAFLD | 1 | USA | Discovery |
| SRR5275440 | F | 29 | 54 | NAFLD | 0 | USA | Discovery |
| SRR5275441 | F | 27 | 40 | NAFLD | 0 | USA | Discovery |
| SRR5275442 | M | 37.1 | 23 | NAFLD | 1 | USA | Discovery |
| SRR5275443 | M | 32 | 62 | NAFLD | 3 | USA | Discovery |
| SRR5275444 | M | 37 | 47 | NAFLD | 1 | USA | Discovery |
| SRR5275445 | M | 23.8 | 44 | NAFLD | 0 | USA | Discovery |
| SRR5275446 | M | 28.7 | 56 | NAFLD | 0 | USA | Discovery |
| SRR5275447 | M | 26.9 | 61 | NAFLD | 0 | USA | Discovery |
| SRR5275448 | F | 35.9 | 22 | NAFLD | 0 | USA | Discovery |
| SRR5275449 | F | 21 | 69 | NAFLD | 4 | USA | Discovery |
| SRR5275450 | F | 25.3 | 22 | NAFLD | 0 | USA | Discovery |
| SRR5275451 | M | 28.9 | 65 | NAFLD | 1 | USA | Discovery |
| SRR5275452 | F | 30.9 | 45 | NAFLD | 1 | USA | Discovery |
| SRR5275453 | M | 27.7 | 45 | NAFLD | 2 | USA | Discovery |
| SRR5275454 | F | 25.4 | 63 | NAFLD | 0 | USA | Discovery |
| SRR5275455 | F | 49.6 | 49 | NAFLD | 2 | USA | Discovery |
| SRR5275456 | F | 30.7 | 68 | NAFLD | 4 | USA | Discovery |
| SRR5275457 | F | 21.4 | 60 | NAFLD | 4 | USA | Discovery |
| SRR5275458 | F | 35.9 | 46 | NAFLD | 1 | USA | Discovery |
| SRR5275459 | M | 29.8 | 69 | NAFLD | 2 | USA | Discovery |
| SRR5275460 | F | 32.8 | 66 | NAFLD | 4 | USA | Discovery |
| SRR5275461 | F | 40.5 | 50 | NAFLD | 1 | USA | Discovery |
| SRR5275462 | M | 32 | 71 | NAFLD | 1 | USA | Discovery |
| SRR5275463 | M | 40.7 | 51 | NAFLD | 1 | USA | Discovery |
| SRR5275464 | F | 28.4 | 61 | NAFLD | 0 | USA | Discovery |
| SRR5275465 | M | 37.9 | 33 | NAFLD | 0 | USA | Discovery |
| SRR5275466 | M | 30.4 | 36 | NAFLD | 2 | USA | Discovery |
| SRR5275467 | F | 30.9 | 66 | NAFLD | 0 | USA | Discovery |
| SRR5275468 | M | 27.6 | 47 | NAFLD | 0 | USA | Discovery |
| SRR5275469 | F | 38.1 | 61 | NAFLD | 1 | USA | Discovery |
| SRR5275470 | M | 39 | 66 | NAFLD | 3 | USA | Discovery |
| SRR5275471 | F | 32.5 | 56 | NAFLD | 0 | USA | Discovery |
| SRR5275472 | F | 31.3 | 37 | NAFLD | 2 | USA | Discovery |
| SRR5275473 | F | 36.6 | 61 | NAFLD | 3 | USA | Discovery |
| SRR5275474 | F | 33.2 | 63 | NAFLD | 4 | USA | Discovery |
| SRR5275475 | F | 28.7 | 39 | NAFLD | 0 | USA | Discovery |
| SRR5275476 | F | 28.3 | 40 | NAFLD | 0 | USA | Discovery |
| SRR5275477 | M | 31 | 64 | NAFLD | 0 | USA | Discovery |
| SRR5275478 | M | 25 | 23 | NAFLD | 0 | USA | Discovery |
| SRR5275479 | F | 19.2 | 49 | NAFLD | 0 | USA | Discovery |
| SRR5275480 | M | 30.9 | 41 | NAFLD | 0 | USA | Discovery |
| SRR5275481 | F | 36.3 | 62 | NAFLD | 4 | USA | Discovery |
| SRR5275482 | M | 29.2 | 65 | NAFLD | 1 | USA | Discovery |
| SRR5275483 | F | 29.3 | 58 | NAFLD | 0 | USA | Discovery |
| SRR5275484 | F | 28.6 | 49 | NAFLD | 0 | USA | Discovery |
| DE-074 | M | 23 | 34 | Control | NA | Germany | Discovery |
| DE-076 | M | 25 | 37 | Control | NA | Germany | Discovery |
| DE-077 | M | 21 | 34 | Control | NA | Germany | Discovery |
| FR-001 | F | 20 | 52 | Control | NA | France | Discovery |
| FR-003 | M | 22 | 70 | Control | NA | France | Discovery |
| FR-024 | F | 18 | 37 | Control | NA | France | Discovery |
| FR-040 | M | 24 | 57 | Control | NA | France | Discovery |
| FR-053 | F | 21 | 62 | Control | NA | France | Discovery |
| FR-110 | F | 25 | 59 | Control | NA | France | Discovery |
| FR-116 | F | 20 | 55 | Control | NA | France | Discovery |
| FR-118 | F | 20 | 64 | Control | NA | France | Discovery |
| FR-121 | M | 25 | 59 | Control | NA | France | Discovery |
| FR-132 | F | 25 | 69 | Control | NA | France | Discovery |
| FR-139 | F | 24 | 61 | Control | NA | France | Discovery |
| FR-142 | F | 25 | 68 | Control | NA | France | Discovery |
| FR-152 | F | 21 | 63 | Control | NA | France | Discovery |
| FR-156 | M | 24 | 62 | Control | NA | France | Discovery |
| FR-169 | F | 24 | 65 | Control | NA | France | Discovery |
| FR-173 | F | 25 | 59 | Control | NA | France | Discovery |
| FR-187 | M | 25 | 70 | Control | NA | France | Discovery |
| FR-192 | M | 24 | 29 | Control | NA | France | Discovery |
| FR-193 | M | 24 | 25 | Control | NA | France | Discovery |
| FR-195 | F | 23 | 63 | Control | NA | France | Discovery |
| FR-198 | M | 25 | 63 | Control | NA | France | Discovery |
| FR-213 | F | 23 | 61 | Control | NA | France | Discovery |
| FR-305 | M | 25 | 50 | Control | NA | France | Discovery |
| FR-316 | M | 23 | 62 | Control | NA | France | Discovery |
| FR-349 | M | 23 | 61 | Control | NA | France | Discovery |
| FR-393 | F | 23 | 67 | Control | NA | France | Discovery |
| FR-399 | F | 22 | 63 | Control | NA | France | Discovery |
| FR-400 | M | 21 | 67 | Control | NA | France | Discovery |
| FR-568 | M | 23 | 35 | Control | NA | France | Discovery |
| FR-684 | M | 25 | 67 | Control | NA | France | Discovery |
| FR-696 | M | 24 | 52 | Control | NA | France | Discovery |
| FR-719 | F | 23 | 49 | Control | NA | France | Discovery |
| FR-721 | F | 20 | 62 | Control | NA | France | Discovery |
| FR-730 | F | 22 | 38 | Control | NA | France | Discovery |
| FR-751 | M | 25 | 66 | Control | NA | France | Discovery |
| MH0006 | F | 22.4 | 59 | Healthy | NA | danish | Validation |
| MH0086 | F | 21.6 | 60 | Healthy | NA | danish | Validation |
| MH0098 | M | 23.9 | 55 | Healthy | NA | danish | Validation |
| MH0113 | M | 23.1 | 55 | Healthy | NA | danish | Validation |
| MH0115 | F | 23.9 | 59 | Healthy | NA | danish | Validation |
| MH0135 | M | 22.6 | 70 | Healthy | NA | danish | Validation |
| MH0149 | F | 22.4 | 50 | Healthy | NA | danish | Validation |
| MH0151 | M | 23.0 | 45 | Healthy | NA | danish | Validation |
| MH0175 | F | 24.5 | 50 | Healthy | NA | danish | Validation |
| MH0179 | F | 24.4 | 60 | Healthy | NA | danish | Validation |
| MH0183 | F | 23.3 | 55 | Healthy | NA | danish | Validation |
| SRR6339475 | NA | NA | NA | NA | NAFLD | USA | Validation |
| SRR6339471 | NA | NA | NA | NA | NAFLD | USA | Validation |
| SRR6339468 | NA | NA | NA | NA | NAFLD | USA | Validation |
| SRR6339462 | NA | NA | NA | NA | NAFLD | USA | Validation |
| SRR6339457 | NA | NA | NA | NA | NAFLD | USA | Validation |
| SRR6339488 | NA | NA | NA | NA | NAFLD | USA | Validation |
| SRR6339495 | NA | NA | NA | NA | NAFLD | USA | Validation |
| SRR6339452 | NA | NA | NA | NA | NAFLD | USA | Validation |
| SRR6339450 | NA | NA | NA | NA | NAFLD | USA | Validation |
| SRR6339485 | NA | NA | NA | NA | NAFLD | USA | Validation |

| **Table S2 The significant differential species between NAFLD and Healthy** | | | | | | | |
| --- | --- | --- | --- | --- | --- | --- | --- |
| **Species ^a^** | **H_mean ^b^** | **N_mean ^c^** | **H_sd ^d^** | **N_sd ^e^** | **lof2 ratio ^f^** | **pvalue ^g^** | **p.adjust ^h^** |
| Bifidobacterium_longum | 1.1281 | 0.8859 | 1.9976 | 3.5042 | -0.3486 | 0.0133 | 0.0347 |
| Adlercreutzia_equolifaciens | 0.0718 | 0.0104 | 0.1414 | 0.0442 | -2.7906 | 0.0002 | 0.0011 |
| Collinsella_aerofaciens | 0.9115 | 0.3823 | 1.0607 | 0.6141 | -1.2534 | 0.0212 | 0.0516 |
| Bacteroides_dorei | 1.7903 | 0.5939 | 3.5703 | 1.6769 | -1.5919 | 0.0000 | 0.0000 |
| Bacteroides_finegoldii | 0.0600 | 0.5016 | 0.1531 | 0.9843 | 3.0643 | 0.0063 | 0.0177 |
| Bacteroides_ovatus | 2.0653 | 4.7042 | 3.2225 | 8.9055 | 1.1876 | 0.0308 | 0.0720 |
| Bacteroides_stercoris | 0.8642 | 4.1655 | 1.8216 | 7.1141 | 2.2690 | 0.0010 | 0.0039 |
| Bacteroidales_bacterium_ph8 | 0.1515 | 0.0652 | 0.2277 | 0.3229 | -1.2164 | 0.0000 | 0.0000 |
| Barnesiella_intestinihominis | 0.7744 | 0.3998 | 1.0334 | 0.9979 | -0.9539 | 0.0002 | 0.0010 |
| Odoribacter_splanchnicus | 0.5867 | 0.1809 | 0.5060 | 0.4347 | -1.6974 | 0.0000 | 0.0000 |
| Parabacteroides_distasonis | 1.3087 | 0.7117 | 2.4723 | 1.8349 | -0.8789 | 0.0053 | 0.0152 |
| Paraprevotella_xylaniphila | 0.0662 | 0.0336 | 0.2891 | 0.1531 | -0.9812 | 0.0197 | 0.0491 |
| Alistipes_finegoldii | 0.2137 | 0.1228 | 0.3873 | 0.3169 | -0.7995 | 0.0025 | 0.0092 |
| Alistipes_indistinctus | 0.1234 | 0.0357 | 0.5403 | 0.1067 | -1.7879 | 0.0003 | 0.0014 |
| Alistipes_putredinis | 0.9153 | 0.6987 | 0.9996 | 1.3588 | -0.3895 | 0.0021 | 0.0079 |
| Alistipes_senegalensis | 0.0439 | 0.0000 | 0.1035 | 0.0000 | NA | 0.0000 | 0.0000 |
| Alistipes_shahii | 0.3005 | 0.1568 | 0.3533 | 0.3063 | -0.9386 | 0.0000 | 0.0002 |
| Lactococcus_lactis | 0.1925 | 0.0240 | 0.4148 | 0.0967 | -3.0037 | 0.0008 | 0.0033 |
| Streptococcus_anginosus | 0.0225 | 0.0119 | 0.0980 | 0.0925 | -0.9143 | 0.0004 | 0.0020 |
| Streptococcus_australis | 0.0261 | 0.0189 | 0.0411 | 0.0450 | -0.4660 | 0.0001 | 0.0006 |
| Streptococcus_thermophilus | 0.7755 | 0.0896 | 2.2937 | 0.2413 | -3.1143 | 0.0011 | 0.0041 |
| Clostridium_bolteae | 0.6099 | 0.6437 | 2.4459 | 1.5724 | 0.0777 | 0.0218 | 0.0521 |
| Flavonifractor_plautii | 0.0196 | 0.0543 | 0.0513 | 0.1388 | 1.4671 | 0.0047 | 0.0140 |
| Eubacterium_eligens | 2.3188 | 0.4986 | 3.1017 | 1.7143 | -2.2176 | 0.0000 | 0.0000 |
| Eubacterium_hallii | 2.0601 | 0.5316 | 2.7180 | 0.8537 | -1.9543 | 0.0000 | 0.0000 |
| Eubacterium_ramulus | 0.4671 | 0.1628 | 0.6385 | 0.3447 | -1.5204 | 0.0001 | 0.0004 |
| Eubacterium_siraeum | 0.6814 | 1.6282 | 1.2036 | 6.7405 | 1.2566 | 0.0005 | 0.0021 |
| Eubacterium_ventriosum | 0.4709 | 0.2605 | 0.5468 | 0.6427 | -0.8544 | 0.0000 | 0.0001 |
| Anaerostipes_hadrus | 0.2136 | 0.0680 | 0.3720 | 0.2426 | -1.6517 | 0.0000 | 0.0000 |
| Ruminococcus_obeum | 1.7214 | 0.5239 | 1.8568 | 1.1477 | -1.7162 | 0.0000 | 0.0000 |
| Ruminococcus_torques | 2.5472 | 2.1543 | 2.1619 | 2.8653 | -0.2417 | 0.0148 | 0.0377 |
| Coprococcus_catus | 0.2567 | 0.1553 | 0.2332 | 0.4550 | -0.7252 | 0.0003 | 0.0015 |
| Dorea_formicigenerans | 0.8557 | 0.4421 | 0.7912 | 0.7835 | -0.9528 | 0.0000 | 0.0001 |
| Dorea_longicatena | 1.3413 | 0.8318 | 1.3325 | 1.4750 | -0.6893 | 0.0027 | 0.0096 |
| Lachnospiraceae_bacterium_1_1_57FAA | 1.4126 | 0.0547 | 4.3049 | 0.1465 | -4.6911 | 0.0000 | 0.0000 |
| Lachnospiraceae_bacterium_5_1_63FAA | 0.3523 | 0.1744 | 0.9995 | 0.4725 | -1.0146 | 0.0032 | 0.0107 |
| Roseburia_inulinivorans | 1.2388 | 0.9409 | 1.1702 | 1.9332 | -0.3968 | 0.0008 | 0.0033 |
| Clostridium_bartlettii | 0.1179 | 0.0312 | 0.1970 | 0.1010 | -1.9184 | 0.0000 | 0.0000 |
| Peptostreptococcaceae_noname_unclassified | 0.1732 | 0.0498 | 0.3454 | 0.2106 | -1.7979 | 0.0050 | 0.0145 |
| Anaerotruncus_unclassified | 0.0302 | 0.0125 | 0.0606 | 0.0367 | -1.2734 | 0.0000 | 0.0003 |
| Faecalibacterium_prausnitzii | 5.8452 | 2.5823 | 4.8547 | 3.7461 | -1.1786 | 0.0000 | 0.0001 |
| Ruminococcus_bromii | 5.5036 | 1.5971 | 6.4013 | 3.4165 | -1.7849 | 0.0000 | 0.0000 |
| Ruminococcus_callidus | 0.2600 | 0.1244 | 0.7459 | 0.4685 | -1.0636 | 0.0001 | 0.0006 |
| Ruminococcus_lactaris | 0.5439 | 0.1819 | 0.9094 | 0.5341 | -1.5802 | 0.0042 | 0.0131 |
| Ruminococcus_sp_5_1_39BFAA | 3.0918 | 1.5653 | 4.1119 | 2.8309 | -0.9820 | 0.0032 | 0.0107 |
| Coprobacillus_unclassified | 0.0718 | 0.1843 | 0.1819 | 0.4103 | 1.3595 | 0.0085 | 0.0225 |
| Dialister_invisus | 0.7315 | 1.3997 | 1.6608 | 3.8972 | 0.9362 | 0.0441 | 0.0973 |
| Parasutterella_excrementihominis | 0.0286 | 0.0228 | 0.0490 | 0.0874 | -0.3221 | 0.0034 | 0.0109 |
| Sutterella_wadsworthensis | 0.3027 | 0.0852 | 0.6046 | 0.3029 | -1.8291 | 0.0039 | 0.0123 |
| Bilophila_unclassified | 0.1484 | 0.1870 | 0.1971 | 0.4641 | 0.3341 | 0.0347 | 0.0796 |
| Bilophila_wadsworthia | 0.0350 | 0.0163 | 0.0774 | 0.0420 | -1.1054 | 0.0078 | 0.0212 |
| Escherichia_coli | 3.1043 | 10.6535 | 6.9588 | 22.2907 | 1.7790 | 0.0364 | 0.0820 |
| Clostridium_clostridioforme | 0.0000 | 0.1592 | 0.0000 | 0.5529 | Inf | 0.0000 | 0.0000 |
| **a: The significant differential species between NAFLD and Health** | | |  |  |  |  |  |
| **b: Average relative abundance of the species in health controls** | | |  |  |  |  |  |
| **c:Average realtive abundance of the species in NAFLD patients** | | |  |  |  |  |  |
| **d:Standard Deviation of the species in health controls** | |  |  |  |  |  |  |
| **e:Standard Deviation of the species in NAFLD patients** | |  |  |  |  |  |  |
| **f:The abundance log2 ratio of the species (NAFLD:Health)** | | |  |  |  |  |  |
| **g: Mann-Whitney U-test** |  |  |  |  |  |  |  |
| **h: Benjamini–Hochberg adjusted p value** |  |  |  |  |  |  |  |

| **Table S3 The significant differential KEGG pathways between NAFLD and Healthy** | | | | | | | |
| --- | --- | --- | --- | --- | --- | --- | --- |
| **KEGG pathways ^a^** | **H_mean ^b^** | **N_mean ^c^** | **H_sd ^d^** | **N_sd ^e^** | **lof2 ratio ^f^** | **pvalue ^g^** | **p.adjust ^h^** |
| UNINTEGRATED | 0.02770 | 0.04513 | 0.00972 | 0.03292 | 0.70441 | 0.00023 | 0.00057 |
| ko00010: Glycolysis / Gluconeogenesis | 0.00069 | 0.00094 | 0.00016 | 0.00042 | 0.44310 | 0.00017 | 0.00048 |
| ko00020: Citrate cycle (TCA cycle) | 0.00048 | 0.00071 | 0.00027 | 0.00044 | 0.57051 | 0.00079 | 0.00144 |
| ko00030: Pentose phosphate pathway | 0.00086 | 0.00120 | 0.00026 | 0.00056 | 0.47961 | 0.00024 | 0.00058 |
| ko00040: Pentose and glucuronate interconversions | 0.00045 | 0.00065 | 0.00018 | 0.00041 | 0.53455 | 0.00921 | 0.01161 |
| ko00051: Fructose and mannose metabolism | 0.00055 | 0.00089 | 0.00019 | 0.00053 | 0.68320 | 0.00002 | 0.00027 |
| ko00052: Galactose metabolism | 0.00049 | 0.00085 | 0.00020 | 0.00057 | 0.78991 | 0.00001 | 0.00025 |
| ko00053: Ascorbate and aldarate metabolism | 0.00013 | 0.00031 | 0.00006 | 0.00039 | 1.18256 | 0.00141 | 0.00230 |
| ko00061: Fatty acid biosynthesis | 0.00063 | 0.00083 | 0.00025 | 0.00035 | 0.38486 | 0.00222 | 0.00315 |
| ko00071: Fatty acid metabolism | 0.00019 | 0.00027 | 0.00005 | 0.00018 | 0.51339 | 0.00552 | 0.00738 |
| ko00121: Secondary bile acid biosynthesis | 0.00031 | 0.00036 | 0.00009 | 0.00013 | 0.24722 | 0.01146 | 0.01429 |
| ko00130: Ubiquinone and other terpenoid-quinone biosynthesis | 0.00012 | 0.00027 | 0.00008 | 0.00027 | 1.16155 | 0.00005 | 0.00031 |
| ko00140: Steroid hormone biosynthesis | 0.00003 | 0.00005 | 0.00002 | 0.00004 | 0.99057 | 0.00036 | 0.00079 |
| ko00190: Oxidative phosphorylation | 0.00022 | 0.00030 | 0.00008 | 0.00012 | 0.47458 | 0.00004 | 0.00031 |
| ko00230: Purine metabolism | 0.00046 | 0.00060 | 0.00011 | 0.00022 | 0.37255 | 0.00019 | 0.00050 |
| ko00240: Pyrimidine metabolism | 0.00057 | 0.00073 | 0.00015 | 0.00023 | 0.36108 | 0.00006 | 0.00031 |
| ko00250: Alanine, aspartate and glutamate metabolism | 0.00105 | 0.00134 | 0.00031 | 0.00042 | 0.35222 | 0.00007 | 0.00031 |
| ko00260: Glycine, serine and threonine metabolism | 0.00055 | 0.00075 | 0.00015 | 0.00033 | 0.46591 | 0.00008 | 0.00032 |
| ko00270: Cysteine and methionine metabolism | 0.00062 | 0.00081 | 0.00017 | 0.00034 | 0.37198 | 0.00099 | 0.00167 |
| ko00280: Valine, leucine and isoleucine degradation | 0.00017 | 0.00025 | 0.00006 | 0.00014 | 0.51942 | 0.00018 | 0.00048 |
| ko00290: Valine, leucine and isoleucine biosynthesis | 0.00141 | 0.00173 | 0.00030 | 0.00056 | 0.30039 | 0.00024 | 0.00058 |
| ko00300: Lysine biosynthesis | 0.00079 | 0.00100 | 0.00025 | 0.00033 | 0.34748 | 0.00026 | 0.00062 |
| ko00310: Lysine degradation | 0.00010 | 0.00019 | 0.00004 | 0.00020 | 0.90172 | 0.00013 | 0.00042 |
| ko00311: Penicillin and cephalosporin biosynthesis | 0.00007 | 0.00016 | 0.00005 | 0.00020 | 1.17295 | 0.00137 | 0.00227 |
| ko00330: Arginine and proline metabolism | 0.00043 | 0.00057 | 0.00013 | 0.00022 | 0.41426 | 0.00018 | 0.00048 |
| ko00340: Histidine metabolism | 0.00064 | 0.00084 | 0.00026 | 0.00032 | 0.38958 | 0.00053 | 0.00105 |
| ko00350: Tyrosine metabolism | 0.00004 | 0.00014 | 0.00008 | 0.00018 | 1.71495 | 0.00312 | 0.00438 |
| ko00360: Phenylalanine metabolism | 0.00013 | 0.00026 | 0.00006 | 0.00023 | 0.99823 | 0.00001 | 0.00025 |
| ko00362: Benzoate degradation | 0.00009 | 0.00011 | 0.00002 | 0.00009 | 0.20791 | 0.03667 | 0.04417 |
| ko00364: Fluorobenzoate degradation | 0.00001 | 0.00005 | 0.00002 | 0.00013 | 3.11326 | 0.00881 | 0.01124 |
| ko00400: Phenylalanine, tyrosine and tryptophan biosynthesis | 0.00051 | 0.00071 | 0.00017 | 0.00029 | 0.49538 | 0.00003 | 0.00027 |
| ko00430: Taurine and hypotaurine metabolism | 0.00016 | 0.00033 | 0.00015 | 0.00026 | 1.01398 | 0.00048 | 0.00100 |
| ko00440: Phosphonate and phosphinate metabolism | 0.00008 | 0.00014 | 0.00004 | 0.00010 | 0.80751 | 0.00167 | 0.00257 |
| ko00450: Selenocompound metabolism | 0.00082 | 0.00100 | 0.00023 | 0.00040 | 0.28954 | 0.00148 | 0.00238 |
| ko00471: D-Glutamine and D-glutamate metabolism | 0.00084 | 0.00123 | 0.00035 | 0.00056 | 0.55689 | 0.00007 | 0.00031 |
| ko00473: D-Alanine metabolism | 0.00058 | 0.00102 | 0.00023 | 0.00068 | 0.79910 | 0.00002 | 0.00026 |
| ko00480: Glutathione metabolism | 0.00027 | 0.00045 | 0.00010 | 0.00035 | 0.69602 | 0.00178 | 0.00271 |
| ko00500: Starch and sucrose metabolism | 0.00048 | 0.00069 | 0.00015 | 0.00032 | 0.52432 | 0.00003 | 0.00027 |
| ko00511: Other glycan degradation | 0.00086 | 0.00139 | 0.00064 | 0.00086 | 0.69687 | 0.00063 | 0.00120 |
| ko00520: Amino sugar and nucleotide sugar metabolism | 0.00055 | 0.00078 | 0.00018 | 0.00032 | 0.51041 | 0.00002 | 0.00027 |
| ko00531: Glycosaminoglycan degradation | 0.00034 | 0.00052 | 0.00028 | 0.00039 | 0.62779 | 0.00625 | 0.00825 |
| ko00540: Lipopolysaccharide biosynthesis | 0.00035 | 0.00078 | 0.00020 | 0.00058 | 1.15546 | 0.00000 | 0.00025 |
| ko00550: Peptidoglycan biosynthesis | 0.00065 | 0.00096 | 0.00025 | 0.00044 | 0.54596 | 0.00008 | 0.00032 |
| ko00561: Glycerolipid metabolism | 0.00016 | 0.00026 | 0.00006 | 0.00021 | 0.66665 | 0.00690 | 0.00900 |
| ko00564: Glycerophospholipid metabolism | 0.00026 | 0.00039 | 0.00007 | 0.00026 | 0.58248 | 0.00162 | 0.00254 |
| ko00620: Pyruvate metabolism | 0.00065 | 0.00087 | 0.00015 | 0.00035 | 0.41554 | 0.00009 | 0.00033 |
| ko00626: Naphthalene degradation | 0.00016 | 0.00022 | 0.00006 | 0.00011 | 0.47069 | 0.00197 | 0.00291 |
| ko00627: Aminobenzoate degradation | 0.00004 | 0.00010 | 0.00003 | 0.00014 | 1.15367 | 0.02471 | 0.03014 |
| ko00630: Glyoxylate and dicarboxylate metabolism | 0.00037 | 0.00062 | 0.00013 | 0.00049 | 0.72431 | 0.00030 | 0.00069 |
| ko00640: Propanoate metabolism | 0.00032 | 0.00046 | 0.00009 | 0.00027 | 0.52512 | 0.00090 | 0.00161 |
| ko00650: Butanoate metabolism | 0.00032 | 0.00048 | 0.00009 | 0.00029 | 0.60592 | 0.00010 | 0.00034 |
| ko00660: C5-Branched dibasic acid metabolism | 0.00089 | 0.00122 | 0.00023 | 0.00065 | 0.46523 | 0.00048 | 0.00100 |
| ko00670: One carbon pool by folate | 0.00082 | 0.00109 | 0.00026 | 0.00038 | 0.41304 | 0.00007 | 0.00031 |
| ko00680: Methane metabolism | 0.00027 | 0.00037 | 0.00008 | 0.00015 | 0.45284 | 0.00007 | 0.00031 |
| ko00710: Carbon fixation in photosynthetic organisms | 0.00095 | 0.00123 | 0.00029 | 0.00053 | 0.37202 | 0.00027 | 0.00062 |
| ko00720: Carbon fixation pathways in prokaryotes | 0.00058 | 0.00079 | 0.00020 | 0.00031 | 0.46236 | 0.00010 | 0.00034 |
| ko00730: Thiamine metabolism | 0.00085 | 0.00106 | 0.00026 | 0.00040 | 0.31610 | 0.00181 | 0.00272 |
| ko00740: Riboflavin metabolism | 0.00042 | 0.00065 | 0.00017 | 0.00038 | 0.65214 | 0.00006 | 0.00031 |
| ko00750: Vitamin B6 metabolism |  | 0.00089 | 0.00019 | 0.00035 | 0.35354 | 0.00099 | 0.00167 |
| ko00760: Nicotinate and nicotinamide metabolism | 0.00046 | 0.00070 | 0.00018 | 0.00037 | 0.59231 | 0.00011 | 0.00035 |
| ko00770: Pantothenate and CoA biosynthesis | 0.00086 | 0.00118 | 0.00023 | 0.00044 | 0.45550 | 0.00001 | 0.00026 |
| ko00780: Biotin metabolism | 0.00052 | 0.00097 | 0.00026 | 0.00072 | 0.89820 | 0.00001 | 0.00025 |
| ko00785: Lipoic acid metabolism | 0.00040 | 0.00086 | 0.00025 | 0.00059 | 1.09914 | 0.00001 | 0.00025 |
| ko00790: Folate biosynthesis | 0.00045 | 0.00073 | 0.00020 | 0.00046 | 0.69785 | 0.00036 | 0.00079 |
| ko00860: Porphyrin and chlorophyll metabolism | 0.00019 | 0.00030 | 0.00009 | 0.00018 | 0.67625 | 0.00014 | 0.00042 |
| ko00900: Terpenoid backbone biosynthesis | 0.00049 | 0.00067 | 0.00017 | 0.00028 | 0.44835 | 0.00016 | 0.00048 |
| ko00908: Zeatin biosynthesis | 0.00026 | 0.00039 | 0.00012 | 0.00018 | 0.59984 | 0.00004 | 0.00031 |
| ko00910: Nitrogen metabolism | 0.00031 | 0.00049 | 0.00011 | 0.00031 | 0.63958 | 0.00014 | 0.00042 |
| ko00920: Sulfur metabolism | 0.00028 | 0.00050 | 0.00026 | 0.00041 | 0.83607 | 0.00214 | 0.00309 |
| ko00970: Aminoacyl-tRNA biosynthesis | 0.00104 | 0.00116 | 0.00021 | 0.00030 | 0.15649 | 0.00878 | 0.01124 |
| ko01040: Biosynthesis of unsaturated fatty acids | 0.00010 | 0.00016 | 0.00005 | 0.00011 | 0.66934 | 0.00064 | 0.00120 |
| ko01051: Biosynthesis of ansamycins | 0.00248 | 0.00318 | 0.00085 | 0.00153 | 0.35719 | 0.00377 | 0.00523 |
| ko01053: Biosynthesis of siderophore group nonribosomal peptides | 0.00004 | 0.00011 | 0.00003 | 0.00017 | 1.36121 | 0.00018 | 0.00048 |
| ko01055: Biosynthesis of vancomycin group antibiotics | 0.00203 | 0.00261 | 0.00058 | 0.00082 | 0.36263 | 0.00005 | 0.00031 |
| ko02010: ABC transporters | 0.00028 | 0.00044 | 0.00010 | 0.00040 | 0.67411 | 0.02110 | 0.02602 |
| ko02020: Two-component system | 0.00012 | 0.00024 | 0.00005 | 0.00027 | 0.97679 | 0.00098 | 0.00167 |
| ko03008: Ribosome biogenesis in eukaryotes | 0.00002 | 0.00003 | 0.00001 | 0.00002 | 0.52410 | 0.00048 | 0.00100 |
| ko03013: RNA transport | 0.00002 | 0.00003 | 0.00001 | 0.00002 | 0.56077 | 0.00206 | 0.00301 |
| ko03018: RNA degradation | 0.00038 | 0.00046 | 0.00009 | 0.00013 | 0.24667 | 0.00070 | 0.00129 |
| ko03020: RNA polymerase | 0.00056 | 0.00064 | 0.00014 | 0.00018 | 0.19055 | 0.00547 | 0.00738 |
| ko03030: DNA replication | 0.00047 | 0.00065 | 0.00016 | 0.00026 | 0.47541 | 0.00006 | 0.00031 |
| ko03060: Protein export | 0.00056 | 0.00078 | 0.00017 | 0.00031 | 0.47425 | 0.00006 | 0.00031 |
| ko03070: Bacterial secretion system | 0.00036 | 0.00055 | 0.00010 | 0.00043 | 0.63342 | 0.00094 | 0.00165 |
| ko03410: Base excision repair | 0.00036 | 0.00052 | 0.00012 | 0.00022 | 0.51335 | 0.00003 | 0.00027 |
| ko03420: Nucleotide excision repair | 0.00038 | 0.00047 | 0.00010 | 0.00014 | 0.27821 | 0.00061 | 0.00118 |
| ko03430: Mismatch repair | 0.00062 | 0.00086 | 0.00021 | 0.00034 | 0.47744 | 0.00007 | 0.00031 |
| ko03440: Homologous recombination | 0.00061 | 0.00080 | 0.00019 | 0.00029 | 0.39277 | 0.00013 | 0.00041 |
| ko04112: Cell cycle - Caulobacter | 0.00074 | 0.00091 | 0.00019 | 0.00027 | 0.28623 | 0.00051 | 0.00102 |
| ko04122: Sulfur relay system | 0.00035 | 0.00057 | 0.00013 | 0.00044 | 0.69574 | 0.00440 | 0.00603 |
| ko04146: Peroxisome | 0.00006 | 0.00004 | 0.00008 | 0.00007 | -0.75923 | 0.03701 | 0.04417 |
| ko04910: Insulin signaling pathway | 0.00007 | 0.00008 | 0.00002 | 0.00003 | 0.28136 | 0.00159 | 0.00253 |
| ko05120: Epithelial cell signaling in Helicobacter pylori infection | 0.00007 | 0.00010 | 0.00002 | 0.00005 | 0.60596 | 0.00003 | 0.00028 |
| **a: The significant differential pathways between NAFLD and Health** | | | | | | | |
| **b: Average relative abundance of the pathways in health controls** | | | | | | | |
| **c:Average realtive abundance of the pathways in NAFLD patients** | | | | | | | |
| **d:Standard Deviation of the pathways in health controls** | | | | | | | |
| **e:Standard Deviation of the pathways in NAFLD patients** | | | | | | | |
| **f:The abundance log2 ratio of the pathways (NAFLD:Health)** | | | | | | | |
| **g: Mann-Whitney U-test** | | | | | | | |
| **h: Benjamini–Hochberg adjusted p value** | | | | | | | |

| **Table S4 Characteristics BA-metabolizing MAGs** | | | | | | | | | | | | | | | | | | | | | | |
| --- | --- | --- | --- | --- | --- | --- | --- | --- | --- | --- | --- | --- | --- | --- | --- | --- | --- | --- | --- | --- | --- | --- |
| **mag ID** | **Scientific name** | **BSH** | **7α-HSDH** | **baiA** | **baiB** | **baiCD** | **baiE** | **baiF** | **baiH** | **baiI** | **#Contigs** | **Contig N50** | **Contig Size** | **Contig Mean** | **shortest Contig** | **Longest Contig** | **#Scaffolds** | **Scaffold N50** | **Scaffold Size** | **Scaffold Mean** | **shortest Scaffold** | **Longest Scaffold** |
| mag_001 | Bacteroides sp. MAG001 | 8 | 18 | 16 | 23 | 9 | 0 | 0 | 20 | 0 | 1908 | 7079 | 5E+06 | 2613 | 104 | 66648 | 1287 | 10083 | 5E+06 | 3827 | 501 | 66648 |
| mag_003 | Bacteroides stercoris MAG003 | 4 | 12 | 10 | 11 | 3 | 0 | 0 | 8 | 0 | 8609 | 2755 | 8E+06 | 952 | 101 | 143093 | 2522 | 7797 | 7E+06 | 2746 | 501 | 143093 |
| mag_005 | Eubacterium rectale MAG005 | 3 | 6 | 5 | 0 | 8 | 0 | 0 | 13 | 0 | 5194 | 11654 | 5E+06 | 973 | 101 | 276911 | 1142 | 31601 | 4E+06 | 3632 | 501 | 400651 |
| mag_006 | Bacteroides thetaiotaomicron MAG006 | 1 | 12 | 12 | 9 | 7 | 0 | 0 | 12 | 0 | 4566 | 23992 | 9E+06 | 2018 | 101 | 190062 | 1622 | 28468 | 9E+06 | 5394 | 508 | 190062 |
| mag_007 | Bacteroides vulgatus MAG007 | 4 | 6 | 6 | 8 | 3 | 0 | 0 | 6 | 0 | 1660 | 7412 | 5E+06 | 2739 | 103 | 48854 | 1165 | 11453 | 5E+06 | 3877 | 503 | 59842 |
| mag_009 | Eubacterium siraeum MAG009 | 1 | 5 | 5 | 2 | 0 | 0 | 0 | 2 | 0 | 2202 | 38856 | 3E+06 | 1585 | 102 | 192760 | 304 | 72945 | 3E+06 | 10143 | 501 | 197018 |
| mag_010 | Ruminococcus bromii MAG010 | 3 | 7 | 7 | 7 | 2 | 0 | 0 | 3 | 0 | 676 | 54940 | 3E+06 | 3770 | 101 | 142265 | 184 | 65604 | 2E+06 | 13250 | 509 | 233974 |
| mag_011 | Roseburia inulinivorans MAG011 | 1 | 3 | 3 | 1 | 8 | 0 | 0 | 14 | 0 | 4908 | 545 | 2E+06 | 440 | 102 | 23723 | 1043 | 1422 | 1E+06 | 1236 | 501 | 23723 |
| mag_012 | Ruminococcus bromii MAG012 | 5 | 13 | 12 | 12 | 0 | 0 | 0 | 6 | 0 | 315 | 89913 | 2E+06 | 7347 | 102 | 182491 | 123 | 110051 | 2E+06 | 18555 | 507 | 219711 |
| mag_013 | Dorea longicatena MAG013 | 1 | 11 | 10 | 0 | 5 | 0 | 3 | 8 | 0 | 1150 | 14334 | 2E+06 | 1673 | 101 | 50840 | 253 | 15747 | 2E+06 | 7021 | 515 | 64295 |
| mag_014 | Bacteroides dorei MAG014 | 1 | 3 | 2 | 3 | 3 | 0 | 0 | 5 | 0 | 1448 | 11724 | 5E+06 | 3553 | 103 | 123058 | 1005 | 18682 | 5E+06 | 5102 | 504 | 147520 |
| mag_018 | Bacteroides vulgatus MAG018 | 4 | 11 | 11 | 7 | 2 | 0 | 0 | 2 | 0 | 6493 | 4569 | 7E+06 | 1070 | 102 | 123185 | 2082 | 13916 | 6E+06 | 2883 | 501 | 123457 |
| mag_021 | Eubacterium rectale MAG021 | 1 | 2 | 2 | 10 | 1 | 0 | 0 | 3 | 0 | 3017 | 455 | 1E+06 | 358 | 101 | 15566 | 475 | 1448 | 601376 | 1266 | 504 | 18655 |
| mag_023 | Ruminococcus bromii MAG023 | 1 | 5 | 3 | 3 | 2 | 0 | 0 | 2 | 0 | 3672 | 3015 | 3E+06 | 782 | 101 | 34176 | 564 | 12774 | 2E+06 | 3913 | 501 | 79980 |
| mag_024 | Eubacterium eligens MAG024 | 1 | 6 | 6 | 2 | 0 | 0 | 0 | 3 | 0 | 2569 | 1085 | 2E+06 | 770 | 151 | 10335 | 1380 | 1298 | 2E+06 | 1171 | 501 | 10335 |
| mag_026 | Bacteroides ovatus MAG026 | 1 | 6 | 6 | 3 | 2 | 0 | 0 | 4 | 0 | 6415 | 4052 | 9E+06 | 1411 | 101 | 94165 | 2338 | 7214 | 8E+06 | 3504 | 501 | 94165 |
| mag_027 | Bacteroides helcogenes MAG027 | 1 | 4 | 3 | 2 | 4 | 0 | 0 | 7 | 0 | 6476 | 2313 | 7E+06 | 1089 | 103 | 27030 | 2927 | 3306 | 6E+06 | 2088 | 501 | 27030 |
| mag_028 | Bacteroides xylanisolvens MAG028 | 2 | 1 | 1 | 4 | 3 | 0 | 0 | 4 | 0 | 7517 | 3982 | 9E+06 | 1150 | 101 | 76662 | 2273 | 7175 | 8E+06 | 3418 | 501 | 76662 |
| mag_030 | Bacteroides vulgatus MAG030 | 2 | 3 | 3 | 4 | 4 | 0 | 0 | 6 | 0 | 6393 | 12648 | 7E+06 | 1032 | 102 | 202507 | 1618 | 23631 | 5E+06 | 3348 | 501 | 202507 |
| mag_031 | Candidatus Arthromitus sp. MAG031 | 2 | 6 | 4 | 7 | 1 | 0 | 0 | 1 | 0 | 178 | 50109 | 1E+06 | 8153 | 104 | 121439 | 75 | 50109 | 1E+06 | 19039 | 514 | 177021 |
| mag_033 | Methanobrevibacter smithii MAG033 | 2 | 1 | 1 | 4 | 0 | 0 | 0 | 2 | 0 | 72 | 101075 | 2E+06 | 24326 | 103 | 311534 | 38 | 127711 | 2E+06 | 45971 | 509 | 424044 |
| mag_035 | Eubacterium siraeum MAG035 | 1 | 3 | 3 | 1 | 0 | 0 | 0 | 0 | 0 | 1943 | 26594 | 3E+06 | 1565 | 102 | 192760 | 271 | 45416 | 3E+06 | 9915 | 502 | 197018 |
| mag_036 | Eubacterium biforme MAG036 | 3 | 1 | 1 | 0 | 5 | 0 | 0 | 7 | 0 | 1226 | 9107 | 2E+06 | 1596 | 101 | 42693 | 267 | 13899 | 2E+06 | 6698 | 501 | 57540 |
| mag_038 | Bacteroides helcogenes MAG038 | 1 | 1 | 1 | 5 | 0 | 0 | 0 | 1 | 0 | 7924 | 9757 | 9E+06 | 1147 | 136 | 103514 | 2239 | 20418 | 8E+06 | 3468 | 501 | 103514 |
| mag_042 | Parabacteroides distasonis MAG042 | 1 | 4 | 3 | 2 | 0 | 0 | 0 | 0 | 0 | 5422 | 34292 | 6E+06 | 1123 | 101 | 488643 | 895 | 48204 | 5E+06 | 5700 | 501 | 488643 |
| mag_043 | Bacteroides plebeius MAG043 | 1 | 3 | 3 | 4 | 2 | 0 | 0 | 5 | 0 | 6058 | 51269 | 6E+06 | 1067 | 102 | 384633 | 1451 | 91668 | 5E+06 | 3664 | 501 | 396943 |
| mag_045 | Bacteroides helcogenes MAG045 | 1 | 5 | 3 | 1 | 0 | 0 | 0 | 1 | 0 | 7683 | 1486 | 6E+06 | 733 | 101 | 34898 | 2100 | 3932 | 5E+06 | 2184 | 501 | 35616 |
| mag_046 | Dorea longicatena MAG046 | 1 | 3 | 3 | 0 | 0 | 0 | 0 | 0 | 0 | 2917 | 3583 | 2E+06 | 799 | 102 | 41577 | 604 | 5380 | 2E+06 | 3001 | 505 | 41577 |
| mag_047 | Bifidobacterium adolescentis MAG047 | 1 | 1 | 0 | 1 | 0 | 0 | 0 | 0 | 0 | 1316 | 26034 | 3E+06 | 1951 | 108 | 110160 | 344 | 33600 | 2E+06 | 6870 | 506 | 110160 |
| mag_049 | Eubacterium_siraeum MAG049 | 1 | 3 | 2 | 1 | 9 | 0 | 0 | 9 | 0 | 1333 | 82188 | 2E+06 | 1860 | 101 | 175138 | 237 | 122314 | 2E+06 | 9509 | 501 | 257393 |
| mag_050 | Coprobacillus sp. MAG050 | 1 | 4 | 4 | 0 | 5 | 0 | 0 | 5 | 0 | 573 | 19622 | 2E+06 | 3666 | 101 | 87796 | 233 | 19780 | 2E+06 | 8694 | 504 | 87796 |
| mag_052 | Ruminococcus bromii MAG052 | 1 | 4 | 3 | 3 | 0 | 0 | 0 | 2 | 0 | 250 | 83271 | 2E+06 | 9097 | 101 | 249491 | 79 | 91429 | 2E+06 | 28387 | 529 | 249491 |
| mag_053 | Eubacterium eligens MAG053 | 1 | 2 | 3 | 2 | 0 | 0 | 0 | 2 | 0 | 3971 | 849 | 2E+06 | 550 | 101 | 42453 | 702 | 7950 | 1E+06 | 2128 | 501 | 46859 |
| mag_054 | Oscillibacter valericigenes MAG054 | 1 | 2 | 2 | 0 | 0 | 0 | 1 | 0 | 0 | 7365 | 1606 | 5E+06 | 722 | 101 | 97965 | 1825 | 4594 | 4E+06 | 2186 | 501 | 97965 |
| mag_056 | Coprococcus sp. MAG056 | 2 | 4 | 4 | 3 | 0 | 0 | 0 | 2 | 0 | 1523 | 971 | 1E+06 | 672 | 109 | 10975 | 442 | 1277 | 525374 | 1189 | 504 | 12219 |
| mag_058 | Ruminococcus bromii MAG058 | 1 | 2 | 2 | 5 | 2 | 0 | 0 | 4 | 0 | 928 | 188328 | 3E+06 | 2914 | 106 | 371462 | 169 | 189644 | 3E+06 | 14936 | 513 | 371462 |
| mag_060 | Eubacterium biforme MAG060 | 1 | 8 | 8 | 0 | 2 | 0 | 0 | 3 | 0 | 2233 | 856 | 1E+06 | 668 | 127 | 6296 | 1180 | 793 | 1E+06 | 1072 | 501 | 6296 |
| mag_061 | Eubacterium rectale MAG061 | 1 | 3 | 2 | 1 | 2 | 0 | 0 | 2 | 0 | 199 | 517 | 83228 | 418 | 103 | 2895 | 35 | 1021 | 33800 | 966 | 510 | 2517 |
| mag_062 | Eubacterium eligens MAG062 | 1 | 8 | 7 | 3 | 2 | 0 | 0 | 3 | 0 | 4662 | 17629 | 4E+06 | 929 | 105 | 337422 | 996 | 38497 | 3E+06 | 3488 | 501 | 337422 |
| mag_063 | Eubacterium eligens MAG063 | 1 | 1 | 1 | 2 | 2 | 0 | 0 | 3 | 0 | 7523 | 8920 | 6E+06 | 761 | 101 | 197061 | 1231 | 39018 | 4E+06 | 3375 | 501 | 197061 |
| mag_064 | Eubacterium eligens MAG064 | 2 | 7 | 7 | 0 | 0 | 0 | 0 | 2 | 0 | 2240 | 8416 | 2E+06 | 1006 | 101 | 50529 | 329 | 14578 | 2E+06 | 5451 | 501 | 61301 |
| mag_071 | Alistipes finegoldii MAG071 | 1 | 2 | 1 | 2 | 1 | 0 | 0 | 1 | 0 | 520 | 131867 | 3E+06 | 6407 | 101 | 349696 | 184 | 136956 | 3E+06 | 17708 | 504 | 349696 |
| mag_074 | Coprococcus sp. MAG074 | 1 | 1 | 1 | 5 |  | 0 | 0 | 1 | 0 | 2478 | 1353 | 2E+06 | 862 | 109 | 10975 | 986 | 2420 | 2E+06 | 1740 | 501 | 13234 |
| mag_076 | Ruminococcus bromii MAG076 | 1 | 2 | 2 | 4 | 2 | 0 | 0 | 5 | 0 | 2856 | 41339 | 3E+06 | 1093 | 101 | 259360 | 422 | 67199 | 3E+06 | 6257 | 501 | 259360 |
| mag_089 | Eubacterium biforme MAG089 | 3 | 1 | 1 | 0 | 0 | 0 | 0 | 2 | 0 | 608 | 18221 | 2E+06 | 3073 | 101 | 46871 | 211 | 19155 | 2E+06 | 8479 | 507 | 46871 |
| mag_097 | Bacteroides ovatus MAG097 | 2 | 2 | 1 | 3 | 0 | 0 | 0 | 0 | 0 | 3059 | 2616 | 3E+06 | 1061 | 101 | 23960 | 1063 | 4679 | 3E+06 | 2630 | 501 | 34946 |
| mag_117 | Bacteroides xylanisolvens MAG117 | 2 | 3 | 3 | 2 | 1 | 0 | 0 | 1 | 0 | 3132 | 2942 | 3E+06 | 1096 | 101 | 27621 | 1035 | 5370 | 3E+06 | 2825 | 501 | 32202 |
| mag_128 | Bacteroides ovatus MAG128 | 1 | 2 | 1 | 2 | 1 | 0 | 0 | 1 | 0 | 4907 | 8173 | 5E+06 | 945 | 110 | 91852 | 1266 | 18114 | 4E+06 | 2944 | 501 | 91852 |
| mag_143 | Mycoplasma sp. MAG143 | 1 | 1 | 1 | 0 | 0 | 0 | 0 | 0 | 0 | 609 | 60907 | 548023 | 900 | 101 | 78319 | 25 | 60907 | 476825 | 19073 | 521 | 78319 |
| mag_198 | Ruminococcus bromii MAG198 | 1 | 1 | 1 | 1 | 1 | 0 | 0 | 1 | 0 | 1154 | 40822 | 2E+06 | 1484 | 102 | 190943 | 151 | 46700 | 1E+06 | 9735 | 504 | 190943 |

| **Table S5 The abundance change of BA-metabolizing MAGs between NAFLD and Healthy** | | | | | | | | | | | | | |
| --- | --- | --- | --- | --- | --- | --- | --- | --- | --- | --- | --- | --- | --- |
| **MAG ID** | **microbial name** | **phylum** | **class** | **order** | **family** | **genus** | **species** | **H_mean** | **N_mean** | **H_sd** | **N_sd** | **pvalue** | **FDR** |
| mag_001 | Bacteroides sp. MAG001 | Bacteroidetes | Bacteroidia | Bacteroidales | Bacteroidaceae | Bacteroides | Bacteroides.sp | 2.1218 | 3.9063 | 1.3362 | 3.6623 | 0.0161 | 0.0336 |
| mag_003 | Bacteroides stercoris MAG003 | Bacteroidetes | Bacteroidia | Bacteroidales | Bacteroidaceae | Bacteroides | Bacteroides stercoris | 0.4086 | 2.2265 | 0.7275 | 3.4019 | 0.0028 | 0.0107 |
| mag_005 | Eubacterium rectale MAG005 | Firmicutes | Clostridia | Clostridiales | Eubacteriaceae | Eubacterium | Eubacterium rectale | 2.0223 | 4.2526 | 1.7630 | 4.4451 | 0.0055 | 0.0137 |
| mag_006 | Bacteroides thetaiotaomicron MAG006 | Bacteroidetes | Bacteroidia | Bacteroidales | Bacteroidaceae | Bacteroides | Bacteroides thetaiotaomicron | 0.2515 | 0.8141 | 0.6347 | 2.6701 | 0.7420 | 0.7894 |
| mag_007 | Bacteroides vulgatus MAG007 | Bacteroidetes | Bacteroidia | Bacteroidales | Bacteroidaceae | Bacteroides | Bacteroides vulgatus | 1.6648 | 2.9960 | 0.9881 | 2.6762 | 0.0138 | 0.0314 |
| mag_009 | Eubacterium siraeum MAG009 | Firmicutes | Clostridia | Clostridiales | Eubacteriaceae | Eubacterium | Eubacterium siraeum | 0.3959 | 1.3480 | 0.8327 | 4.9658 | 0.1144 | 0.1733 |
| mag_010 | Ruminococcus bromii MAG010 | Firmicutes | Clostridia | Clostridiales | Ruminococcaceae | Ruminococcus | Ruminococcus bromii | 2.1374 | 1.5080 | 6.6313 | 8.5019 | 0.0033 | 0.0119 |
| mag_011 | Roseburia inulinivorans MAG011 | Firmicutes | Clostridia | Clostridiales | Lachnospiraceae | Roseburia | Roseburia_inulinivorans | 1.5529 | 0.5485 | 3.1677 | 1.9142 | 0.0003 | 0.0018 |
| mag_012 | Ruminococcus bromii MAG012 | Firmicutes | Clostridia | Clostridiales | Ruminococcaceae | Ruminococcus | Ruminococcus bromii | 3.3496 | 1.8611 | 4.0625 | 3.9114 | 0.0013 | 0.0063 |
| mag_013 | Dorea longicatena MAG013 | Firmicutes | Clostridia | Clostridiales | Lachnospiraceae | Dorea | Dorea longicatena | 1.9465 | 1.1824 | 2.3434 | 1.6881 | 0.0044 | 0.0119 |
| mag_014 | Bacteroides dorei MAG014 | Bacteroidetes | Bacteroidia | Bacteroidales | Bacteroidaceae | Bacteroides | Bacteroides dorei | 1.5067 | 0.8293 | 3.1250 | 2.6227 | 0.0014 | 0.0063 |
| mag_018 | Bacteroides vulgatus MAG018 | Bacteroidetes | Bacteroidia | Bacteroidales | Bacteroidaceae | Bacteroides | Bacteroides vulgatus | 0.7196 | 1.2005 | 1.0258 | 2.4133 | 0.9128 | 0.9315 |
| mag_021 | Eubacterium rectale MAG021 | Firmicutes | Clostridia | Clostridiales | Eubacteriaceae | Eubacterium | Eubacterium rectale | 1.3200 | 0.6215 | 2.4874 | 2.0788 | 0.0159 | 0.0336 |
| mag_023 | Ruminococcus bromii MAG023 | Firmicutes | Clostridia | Clostridiales | Ruminococcaceae | Ruminococcus | Ruminococcus bromii | 1.3600 | 0.9088 | 1.8616 | 2.6412 | 0.0003 | 0.0018 |
| mag_024 | Eubacterium eligens MAG024 | Firmicutes | Clostridia | Clostridiales | Eubacteriaceae | Eubacterium | Eubacterium eligens | 3.5038 | 0.0000 | 8.3930 | 0.0000 | 0.0000 | 0.0000 |
| mag_026 | Bacteroides ovatus MAG026 | Bacteroidetes | Bacteroidia | Bacteroidales | Bacteroidaceae | Bacteroides | Bacteroides ovatus | 1.1164 | 2.2274 | 1.0279 | 2.6345 | 0.0037 | 0.0119 |
| mag_027 | Bacteroides helcogenes MAG027 | Bacteroidetes | Bacteroidia | Bacteroidales | Bacteroidaceae | Bacteroides | Bacteroides helcogenes | 0.4710 | 0.9755 | 0.9803 | 2.8098 | 0.6096 | 0.6927 |
| mag_028 | Bacteroides xylanisolvens MAG028 | Bacteroidetes | Bacteroidia | Bacteroidales | Bacteroidaceae | Bacteroides | Bacteroides xylanisolvens | 0.7569 | 1.3416 | 1.5448 | 2.4727 | 0.6082 | 0.6927 |
| mag_030 | Bacteroides vulgatus MAG030 | Bacteroidetes | Bacteroidia | Bacteroidales | Bacteroidaceae | Bacteroides | Bacteroides vulgatus | 2.3593 | 3.3305 | 2.0397 | 2.7624 | 0.0295 | 0.0547 |
| mag_031 | Candidatus Arthromitus sp. MAG031 | Firmicutes | Clostridia | Clostridiales | Clostridiaceae | Candidatus Arthromitus | Candidatus Arthromitus sp. | 1.9276 | 0.7861 | 7.5841 | 3.7133 | 0.9705 | 0.9705 |
| mag_033 | Methanobrevibacter smithii MAG033 | Euryarchaeota | Methanobacteria | Methanobacteriales | Methanobacteriaceae | Methanobrevibacter | Methanobrevibacter smithii | 2.1799 | 0.6421 | 6.1765 | 2.3282 | 0.0062 | 0.0147 |
| mag_035 | Eubacterium siraeum MAG035 | Firmicutes | Clostridia | Clostridiales | Eubacteriaceae | Eubacterium | Eubacterium siraeum | 0.3468 | 1.0903 | 0.7475 | 4.9932 | 0.0702 | 0.1188 |
| mag_036 | Eubacterium biforme MAG036 | Firmicutes | Erysipelotrichia | Erysipelotrichales | Erysipelotrichaceae | Erysipelotrichaceae_noname | Eubacterium biforme | 0.6975 | 1.2727 | 1.1826 | 5.9292 | 0.0255 | 0.0490 |
| mag_038 | Bacteroides helcogenes MAG038 | Bacteroidetes | Bacteroidia | Bacteroidales | Bacteroidaceae | Bacteroides | Bacteroides helcogenes | 0.9460 | 0.8548 | 2.1222 | 2.1895 | 0.2003 | 0.2635 |
| mag_042 | Parabacteroides distasonis MAG042 | Bacteroidetes | Bacteroidia | Bacteroidales | Porphyromonadaceae | Parabacteroides | Parabacteroides distasonis | 1.3568 | 1.6233 | 1.1420 | 2.3602 | 0.4703 | 0.5736 |
| mag_043 | Bacteroides plebeius MAG043 | Bacteroidetes | Bacteroidia | Bacteroidales | Bacteroidaceae | Bacteroides | Bacteroides plebeius | 1.4637 | 1.4299 | 4.1699 | 7.9144 | 0.0925 | 0.1492 |
| mag_045 | Bacteroides helcogenes MAG045 | Bacteroidetes | Bacteroidia | Bacteroidales | Bacteroidaceae | Bacteroides | Bacteroides helcogenes | 0.9160 | 0.4873 | 1.7217 | 2.6683 | 0.0000 | 0.0000 |
| mag_046 | Dorea longicatena MAG046 | Firmicutes | Clostridia | Clostridiales | Lachnospiraceae | Dorea | Dorea longicatena | 0.6541 | 1.0702 | 0.6968 | 2.3744 | 0.6047 | 0.6927 |
| mag_047 | Bifidobacterium adolescentis MAG047 | Actinobacteria | Actinobacteria | Bifidobacteriales | Bifidobacteriaceae | Bifidobacterium | Bifidobacterium adolescentis | 1.0769 | 1.3547 | 1.6869 | 2.9197 | 0.7264 | 0.7894 |
| mag_049 | Eubacterium_siraeum MAG049 | Firmicutes | Clostridia | Clostridiales | Eubacteriaceae | Eubacterium | Eubacterium siraeum | 1.1998 | 2.3082 | 2.6203 | 5.5238 | 0.6800 | 0.7555 |
| mag_050 | Coprobacillus sp. MAG050 | Firmicutes | Erysipelotrichia | Erysipelotrichales | Erysipelotrichaceae | Coprobacillus | Coprobacillus sp. | 0.5055 | 1.4094 | 1.9069 | 7.2900 | 0.7663 | 0.7983 |
| mag_052 | Ruminococcus bromii MAG052 | Firmicutes | Clostridia | Clostridiales | Ruminococcaceae | Ruminococcus | Ruminococcus bromii | 1.8591 | 0.2546 | 6.1873 | 1.4143 | 0.0241 | 0.0482 |
| mag_053 | Eubacterium eligens MAG053 | Firmicutes | Clostridia | Clostridiales | Eubacteriaceae | Eubacterium | Eubacterium eligens | 1.2675 | 0.2156 | 2.8228 | 1.4281 | 0.0012 | 0.0063 |
| mag_054 | Oscillibacter valericigenes MAG054 | Firmicutes | Clostridia | Clostridiales | Oscillospiraceae | Oscillibacter | Oscillibacter valericigenes | 0.8359 | 0.6511 | 1.5829 | 2.2770 | 0.0021 | 0.0086 |
| mag_056 | Coprococcus sp. MAG056 | Firmicutes | Clostridia | Clostridiales | Lachnospiraceae | Coprococcus | Coprococcus sp. | 2.1944 | 0.1346 | 5.0385 | 1.0691 | 0.0000 | 0.0001 |
| mag_058 | Ruminococcus bromii MAG058 | Firmicutes | Clostridia | Clostridiales | Ruminococcaceae | Ruminococcus | Ruminococcus bromii | 0.3856 | 0.9121 | 0.9096 | 5.4415 | 0.0045 | 0.0119 |
| mag_060 | Eubacterium biforme MAG060 | Firmicutes | Erysipelotrichia | Erysipelotrichales | Erysipelotrichaceae | Erysipelotrichaceae_noname | Eubacterium biforme | 1.5850 | 0.0000 | 3.6822 | 0.0000 | 0.0000 | 0.0000 |
| mag_061 | Eubacterium rectale MAG061 | Firmicutes | Clostridia | Clostridiales | Eubacteriaceae | Eubacterium | Eubacterium rectale | 1.5568 | 0.0000 | 3.5333 | 0.0000 | 0.0000 | 0.0003 |
| mag_062 | Eubacterium eligens MAG062 | Firmicutes | Clostridia | Clostridiales | Eubacteriaceae | Eubacterium | Eubacterium eligens | 2.4654 | 1.1668 | 2.3986 | 3.4539 | 0.0000 | 0.0000 |
| mag_063 | Eubacterium eligens MAG063 | Firmicutes | Clostridia | Clostridiales | Eubacteriaceae | Eubacterium | Eubacterium eligens | 0.7794 | 0.3268 | 2.3797 | 1.5755 | 0.1564 | 0.2234 |
| mag_064 | Eubacterium eligens MAG064 | Firmicutes | Clostridia | Clostridiales | Eubacteriaceae | Eubacterium | Eubacterium eligens | 1.1465 | 0.8756 | 1.9296 | 2.1022 | 0.3537 | 0.4535 |
| mag_071 | Alistipes finegoldii MAG071 | Bacteroidetes | Bacteroidia | Bacteroidales | Rikenellaceae | Alistipes | Alistipes finegoldii | 0.8159 | 0.9456 | 1.1706 | 2.3695 | 0.0713 | 0.1188 |
| mag_074 | Coprococcus sp. MAG074 | Firmicutes | Clostridia | Clostridiales | Lachnospiraceae | Coprococcus | Coprococcus sp. | 1.4020 | 0.1333 | 4.7778 | 1.0555 | 0.0042 | 0.0119 |
| mag_076 | Ruminococcus bromii MAG076 | Firmicutes | Clostridia | Clostridiales | Ruminococcaceae | Ruminococcus | Ruminococcus bromii | 1.1882 | 0.2236 | 4.2369 | 1.1679 | 0.1424 | 0.2094 |
| mag_089 | Eubacterium biforme MAG089 | Firmicutes | Erysipelotrichia | Erysipelotrichales | Erysipelotrichaceae | Erysipelotrichaceae_noname | Eubacterium biforme | 0.8031 | 1.1905 | 1.3605 | 4.9693 | 0.0458 | 0.0818 |
| mag_097 | Bacteroides ovatus MAG097 | Bacteroidetes | Bacteroidia | Bacteroidales | Bacteroidaceae | Bacteroides | Bacteroides ovatus | 0.3837 | 1.2308 | 0.6059 | 2.5237 | 0.1656 | 0.2237 |
| mag_117 | Bacteroides xylanisolvens MAG117 | Bacteroidetes | Bacteroidia | Bacteroidales | Bacteroidaceae | Bacteroides | Bacteroides xylanisolvens | 0.9466 | 2.0396 | 0.7765 | 2.4522 | 0.0041 | 0.0119 |
| mag_128 | Bacteroides ovatus MAG128 | Bacteroidetes | Bacteroidia | Bacteroidales | Bacteroidaceae | Bacteroides | Bacteroides ovatus | 0.9428 | 0.6461 | 2.5624 | 2.3693 | 0.0980 | 0.1531 |
| mag_143 | Mycoplasma sp. MAG143 | Tenericutes | Mollicutes | Mycoplasmatales | Mycoplasmataceae | Mycoplasma | Mycoplasma sp. | 1.2968 | 0.1173 | 6.2349 | 1.0620 | 0.1641 | 0.2237 |
| mag_198 | Ruminococcus bromii MAG198 | Firmicutes | Clostridia | Clostridiales | Ruminococcaceae | Ruminococcus | Ruminococcus bromii | 0.3791 | 0.8034 | 1.0480 | 2.2893 | 0.4575 | 0.5719 |
|  | signifcantly increased in NAFLD | | |  |  |  |  |  |  |  |  |  |  |
|  | signifcantly decreased in NAFLD | |  |  |  |  |  |  |  |  |  |  |  |

| **Table S6 Modularity analysis of BA-metabolizing community in healthy and NAFLD** | | | |
| --- | --- | --- | --- |
| **Healthy** |  | **NAFLD** |  |
| **Moudle ID** | **HQMGs in moudle** | **Moudle ID** | **HQMGs in moudle** |
| **module 0** | Bacteroides ovatus MGS026 | **module 0** | Ruminococcus bromii MGS010 |
|  | Eubacterium eligens MGS062 |  | Methanobrevibacter smithii MGS033 |
|  | Bacteroides xylanisolvens MGS117 |  | Eubacterium biforme MGS036 |
| **module 1** | Bacteroides sp. MGS001 |  | Bacteroides helcogenes MGS045 |
|  | Bacteroides vulgatus MGS007 |  | Bifidobacterium adolescentis MGS047 |
|  | Dorea longicatena MGS013 |  | Coprobacillus sp. MGS050 |
|  | Bacteroides helcogenes MGS027 |  | Eubacterium eligens MGS053 |
|  | Bacteroides vulgatus MGS030 |  | Ruminococcus bromii MGS058 |
|  | Bacteroides helcogenes MGS038 |  | Ruminococcus bromii MGS076 |
|  | Parabacteroides distasonis MGS042 |  | Eubacterium biforme MGS089 |
| **module 2** | Bacteroides xylanisolvens MGS028 |  | Mycoplasma sp. MGS143 |
|  | Bacteroides ovatus MGS097 | **module 1** | Ruminococcus bromii MGS012 |
| **module 3** | Eubacterium siraeum MGS009 |  | Eubacterium rectale MGS021 |
|  | Eubacterium siraeum MGS035 |  | Ruminococcus bromii MGS023 |
|  | Dorea longicatena MGS046 |  | Candidatus Arthromitus sp. MGS031 |
|  | Ruminococcus bromii MGS010 |  | Eubacterium eligens MGS062 |
| **module 4** | Bacteroides thetaiotaomicron MGS006 | **module 2** | Roseburia inulinivorans MGS011 |
|  | Bacteroides plebeius MGS043 |  | Ruminococcus bromii MGS198 |
|  | Bacteroides ovatus MGS128 | **module 3** | Bacteroides ovatus MGS026 |
|  | Mycoplasma sp. MGS143 |  | Bacteroides xylanisolvens MGS117 |
| **module 5** | Candidatus Arthromitus sp. MGS031 | **module 4** | Bacteroides xylanisolvens MGS028 |
|  | Bacteroides helcogenes MGS045 |  | Bacteroides ovatus MGS097 |
|  | Ruminococcus bromii MGS052 | **module 5** | Bacteroides sp. MGS001 |
| **module 6** | Eubacterium rectale MGS021 |  | Bacteroides stercoris MGS003 |
|  | Oscillibacter valericigenes MGS054 |  | Bacteroides vulgatus MGS007 |
|  | Eubacterium biforme MGS060 |  | Bacteroides vulgatus MGS018 |
|  | Bacteroides vulgatus MGS018 |  | Bacteroides dorei MGS014 |
|  | Ruminococcus bromii MGS198 |  | Bacteroides vulgatus MGS030 |
| **module 7** | Eubacterium biforme MGS036 |  | Parabacteroides distasonis MGS042 |
|  | Eubacterium biforme MGS089 |  | Dorea longicatena MGS046 |
| **module 8** | Bifidobacterium adolescentis MGS047 |  | Oscillibacter valericigenes MGS054 |
|  | Coprobacillus sp. MGS050 | **module 6** | Eubacterium_siraeum MGS049 |
|  | Eubacterium eligens MGS063 |  | Bacteroides plebeius MGS043 |
|  | Alistipes finegoldii MGS071 |  | Ruminococcus bromii MGS052 |
|  | Ruminococcus bromii MGS076 |  | Bacteroides ovatus MGS128 |
| **module 9** | Eubacterium eligens MGS053 | **module 7** | Eubacterium siraeum MGS009 |
|  | Coprococcus sp. MGS056 |  | Eubacterium siraeum MGS035 |
|  | Coprococcus sp. MGS074 | **module 8** | Dorea longicatena MGS013 |
| **module 10** | Ruminococcus bromii MGS012 |  | Coprococcus sp. MGS056 |
|  | Bacteroides dorei MGS014 |  | Coprococcus sp. MGS074 |
| **module 11** | Eubacterium rectale MGS005 | **module 9** | Eubacterium eligens MGS024 |
| **module 12** | Roseburia inulinivorans MGS011 |  | Bacteroides helcogenes MGS038 |
|  | Ruminococcus bromii MGS023 |  | Eubacterium eligens MGS063 |
|  | Methanobrevibacter smithii MGS033 |  | Alistipes finegoldii MGS071 |
|  | Ruminococcus bromii MGS058 | **module 10** | Eubacterium rectale MGS005 |
|  | Eubacterium_siraeum MGS049 | **module 11** | Eubacterium eligens MGS064 |
|  | Eubacterium eligens MGS024 |  |  |

| **Table S7 Differential species among three group** | | | | | | | | | |
| --- | --- | --- | --- | --- | --- | --- | --- | --- | --- |
| **Species** | **Health_mean** | **Normal-BA_mean** | **High-BA_mean** | **Health_sd** | **Normal-BA_sd** | **High-BA_sd** | **FDR(High-BA vs Health)** | **FDR(Normal-BA vs Health)** | **FDR(High-BA vs Normal-BA)** |
| Flavonifractor plautii | 0.25641 | 0.58596 | 0.35902 | 0.38041 | 0.47689 | 0.35759 | 0.39857 | 0.00238 | 0.06431 |
| Ruminococcus bromii | 1.89002 | 0.62686 | 1.39507 | 1.21155 | 1.05165 | 1.18968 | 0.04716 | 0.00000 | 0.00485 |
| Escherichia coli | 1.55307 | 1.47886 | 2.51532 | 1.05725 | 1.05511 | 1.17339 | 0.00145 | 1.00000 | 0.00036 |
| * FDR: Dunn tests adjusted by Benjamini–Hochberg | | | | | | | | | |

| **Table S8 Differential EC among three group** | | | | | | | | | |
| --- | --- | --- | --- | --- | --- | --- | --- | --- | --- |
| **EC** | **Health_mean** | **Normal-BA_mean** | **High-BA_mean** | **Health_sd** | **Normal -BA_sd** | **High-BA_sd** | **FDR(High-BA vs Health)** | **FDR(Normal-BA vs Health)** | **FDR(High-BA vs Normal-BA)** |
| ***1.1.1.1: Alcohol dehydrogenase*** | 0.007 | 0.004 | 0.007 | 0.003 | 0.004 | 0.005 | 0.064 | 0.000 | 0.011 |
| ***1.1.1.169: 2-dehydropantoate 2-reductase*** | 0.012 | 0.011 | 0.009 | 0.003 | 0.004 | 0.003 | 0.000 | 0.101 | 0.003 |
| ***1.1.1.262: 4-hydroxythreonine-4-phosphate dehydrogenase*** | 0.006 | 0.009 | 0.007 | 0.003 | 0.004 | 0.003 | 0.124 | 0.000 | 0.030 |
| ***1.1.1.38: Malate dehydrogenase*** | 0.003 | 0.001 | 0.003 | 0.001 | 0.001 | 0.003 | 0.104 | 0.000 | 0.003 |
| ***1.1.1.6: Glycerol dehydrogenase*** | 0.002 | 0.001 | 0.003 | 0.002 | 0.001 | 0.002 | 0.880 | 0.000 | 0.000 |
| ***1.1.1.79: Glyoxylate reductase (NADP(+))*** | 0.000 | 0.000 | 0.003 | 0.001 | 0.000 | 0.004 | 0.000 | 1.000 | 0.000 |
| ***1.1.3.15: (S)-2-hydroxy-acid oxidase*** | 0.002 | 0.001 | 0.002 | 0.002 | 0.002 | 0.001 | 0.429 | 0.000 | 0.016 |
| ***1.12.99.6: Hydrogenase (acceptor)*** | 0.001 | 0.001 | 0.005 | 0.001 | 0.001 | 0.007 | 0.000 | 0.950 | 0.001 |
| ***1.17.1.4: Xanthine dehydrogenase*** | 0.002 | 0.001 | 0.003 | 0.001 | 0.000 | 0.003 | 0.238 | 0.000 | 0.000 |
| ***1.17.1.8: 4-hydroxy-tetrahydrodipicolinate reductase*** | 0.019 | 0.017 | 0.013 | 0.003 | 0.004 | 0.006 | 0.000 | 0.035 | 0.034 |
| ***1.18.1.2: Ferredoxin--NADP(+) reductase*** | 0.004 | 0.002 | 0.003 | 0.002 | 0.002 | 0.002 | 0.011 | 0.000 | 0.025 |
| ***1.20.4.1: Arsenate reductase (glutaredoxin)*** | 0.002 | 0.001 | 0.003 | 0.001 | 0.001 | 0.004 | 1.000 | 0.000 | 0.000 |
| ***1.3.1.12: Prephenate dehydrogenase*** | 0.004 | 0.003 | 0.002 | 0.002 | 0.003 | 0.001 | 0.000 | 0.001 | 0.042 |
| ***2.7.2.11: Glutamate 5-kinase*** | 0.016 | 0.015 | 0.013 | 0.004 | 0.003 | 0.004 | 0.000 | 0.115 | 0.044 |
| ***2.7.6.2: Thiamine diphosphokinase*** | 0.003 | 0.003 | 0.002 | 0.001 | 0.002 | 0.001 | 0.000 | 0.025 | 0.038 |
| ***2.7.7.22: Mannose-1-phosphate guanylyltransferase (GDP)*** | 0.001 | 0.003 | 0.001 | 0.002 | 0.004 | 0.001 | 0.190 | 0.000 | 0.028 |
| ***2.7.7.24: Glucose-1-phosphate thymidylyltransferase*** | 0.019 | 0.018 | 0.016 | 0.004 | 0.004 | 0.005 | 0.001 | 0.520 | 0.011 |
| ***2.7.7.65: Diguanylate cyclase*** | 0.001 | 0.001 | 0.007 | 0.002 | 0.001 | 0.009 | 0.000 | 1.000 | 0.000 |
| ***2.7.7.77: Molybdenum cofactor guanylyltransferase*** | 0.001 | 0.001 | 0.002 | 0.001 | 0.001 | 0.002 | 1.000 | 0.001 | 0.000 |
| ***2.7.8.13: Phospho-N-acetylmuramoyl-pentapeptide-transferase*** | 0.018 | 0.016 | 0.014 | 0.003 | 0.004 | 0.004 | 0.000 | 0.031 | 0.019 |
| ***2.7.8.37: Alpha-D-ribose 1-methylphosphonate 5-triphosphate synthase*** | 0.001 | 0.000 | 0.004 | 0.001 | 0.001 | 0.006 | 0.001 | 1.000 | 0.000 |
| ***2.8.1.1: Thiosulfate sulfurtransferase*** | 0.001 | 0.001 | 0.005 | 0.001 | 0.001 | 0.007 | 0.001 | 1.000 | 0.001 |
| ***2.8.1.10: Thiazole synthase*** | 0.009 | 0.007 | 0.005 | 0.003 | 0.003 | 0.002 | 0.000 | 0.003 | 0.004 |
| ***3.1.1.11: Pectinesterase*** | 0.006 | 0.014 | 0.007 | 0.004 | 0.011 | 0.006 | 1.000 | 0.001 | 0.005 |
| ***3.1.2.12: S-formylglutathione hydrolase*** | 0.000 | 0.000 | 0.003 | 0.001 | 0.000 | 0.005 | 0.000 | 1.000 | 0.001 |
| ***3.1.26.11: Ribonuclease Z*** | 0.013 | 0.012 | 0.009 | 0.003 | 0.004 | 0.004 | 0.000 | 0.322 | 0.003 |
| ***3.1.3.10: Glucose-1-phosphatase*** | 0.000 | 0.000 | 0.004 | 0.001 | 0.000 | 0.006 | 0.000 | 1.000 | 0.001 |
| ***3.1.3.23: Sugar-phosphatase*** | 0.001 | 0.000 | 0.005 | 0.001 | 0.001 | 0.007 | 0.000 | 1.000 | 0.000 |
| ***3.1.3.27: Phosphatidylglycerophosphatase*** | 0.001 | 0.001 | 0.005 | 0.001 | 0.002 | 0.006 | 0.001 | 0.842 | 0.004 |
| ***3.1.3.3: Phosphoserine phosphatase*** | 0.002 | 0.001 | 0.001 | 0.001 | 0.000 | 0.001 | 0.040 | 0.000 | 0.000 |
| ***3.1.3.73: Adenosylcobalamin/alpha-ribazole phosphatase*** | 0.003 | 0.002 | 0.003 | 0.002 | 0.002 | 0.002 | 0.858 | 0.008 | 0.040 |
| ***3.1.3.74: Pyridoxal phosphatase*** | 0.001 | 0.000 | 0.004 | 0.001 | 0.001 | 0.005 | 0.001 | 1.000 | 0.000 |
| ***3.1.4.52: Cyclic-guanylate-specific phosphodiesterase*** | 0.001 | 0.001 | 0.008 | 0.002 | 0.001 | 0.011 | 0.001 | 1.000 | 0.000 |
| ***3.2.1.1: Alpha-amylase*** | 0.009 | 0.005 | 0.008 | 0.005 | 0.004 | 0.006 | 0.099 | 0.000 | 0.006 |
| ***3.2.1.28: Alpha,alpha-trehalase*** | 0.000 | 0.000 | 0.003 | 0.001 | 0.000 | 0.004 | 0.000 | 1.000 | 0.000 |
| ***3.4.11.1: Leucyl aminopeptidase*** | 0.000 | 0.000 | 0.003 | 0.001 | 0.000 | 0.004 | 0.000 | 0.971 | 0.001 |
| ***3.4.11.18: Methionyl aminopeptidase*** | 0.035 | 0.031 | 0.026 | 0.005 | 0.006 | 0.008 | 0.000 | 0.006 | 0.049 |
| ***3.4.16.4: Serine-type D-Ala-D-Ala carboxypeptidase*** | 0.011 | 0.007 | 0.012 | 0.005 | 0.005 | 0.008 | 1.000 | 0.000 | 0.000 |
| ***3.4.17.13: Muramoyltetrapeptide carboxypeptidase*** | 0.000 | 0.000 | 0.003 | 0.001 | 0.001 | 0.004 | 0.000 | 0.805 | 0.001 |
| ***3.4.21.88: Repressor LexA*** | 0.015 | 0.009 | 0.013 | 0.004 | 0.005 | 0.006 | 0.011 | 0.000 | 0.011 |
| ***3.5.1.18: Succinyl-diaminopimelate desuccinylase*** | 0.003 | 0.001 | 0.003 | 0.002 | 0.001 | 0.002 | 0.958 | 0.000 | 0.000 |
| ***3.5.1.28: N-acetylmuramoyl-L-alanine amidase*** | 0.008 | 0.006 | 0.011 | 0.003 | 0.004 | 0.009 | 0.992 | 0.004 | 0.016 |
| ***3.5.3.11: Agmatinase*** | 0.003 | 0.001 | 0.003 | 0.001 | 0.001 | 0.002 | 1.000 | 0.000 | 0.000 |
| ***3.5.4.2: Adenine deaminase*** | 0.009 | 0.005 | 0.007 | 0.004 | 0.003 | 0.003 | 0.091 | 0.000 | 0.004 |
| ***3.5.4.25: GTP cyclohydrolase II*** | 0.001 | 0.001 | 0.002 | 0.001 | 0.001 | 0.002 | 0.725 | 0.002 | 0.014 |
| ***3.5.4.5: Cytidine deaminase*** | 0.003 | 0.002 | 0.002 | 0.001 | 0.001 | 0.002 | 0.135 | 0.000 | 0.021 |
| ***3.6.1.1: Inorganic diphosphatase*** | 0.010 | 0.007 | 0.009 | 0.003 | 0.005 | 0.004 | 0.328 | 0.000 | 0.001 |
| ***3.6.1.13: ADP-ribose diphosphatase*** | 0.003 | 0.002 | 0.003 | 0.001 | 0.002 | 0.002 | 0.074 | 0.000 | 0.017 |
| ***3.6.3.17: Monosaccharide-transporting ATPase*** | 0.011 | 0.007 | 0.013 | 0.004 | 0.004 | 0.009 | 0.909 | 0.000 | 0.000 |
| ***3.6.3.25: Sulfate-transporting ATPase*** | 0.007 | 0.003 | 0.004 | 0.002 | 0.002 | 0.002 | 0.000 | 0.000 | 0.022 |
| ***3.6.3.28: Phosphonate-transporting ATPase*** | 0.004 | 0.001 | 0.002 | 0.002 | 0.001 | 0.002 | 0.000 | 0.000 | 0.018 |
| ***3.6.3.3: Cadmium-exporting ATPase*** | 0.003 | 0.003 | 0.002 | 0.001 | 0.002 | 0.001 | 0.000 | 0.085 | 0.002 |
| ***3.6.3.34: Iron-chelate-transporting ATPase*** | 0.004 | 0.002 | 0.003 | 0.002 | 0.002 | 0.001 | 0.080 | 0.000 | 0.023 |
| ***3.6.3.44: Xenobiotic-transporting ATPase*** | 0.003 | 0.001 | 0.004 | 0.002 | 0.001 | 0.004 | 0.486 | 0.000 | 0.003 |
| ***4.4.1.5: Lactoylglutathione lyase*** | 0.006 | 0.004 | 0.005 | 0.003 | 0.003 | 0.002 | 0.019 | 0.000 | 0.036 |
| ***5.1.3.1: Ribulose-phosphate 3-epimerase*** | 0.018 | 0.017 | 0.015 | 0.003 | 0.004 | 0.004 | 0.000 | 0.042 | 0.037 |
| ***5.1.3.20: ADP-glyceromanno-heptose 6-epimerase*** | 0.001 | 0.002 | 0.003 | 0.001 | 0.003 | 0.003 | 0.000 | 0.509 | 0.002 |
| ***5.1.3.4: L-ribulose-5-phosphate 4-epimerase*** | 0.004 | 0.003 | 0.005 | 0.003 | 0.003 | 0.005 | 0.964 | 0.004 | 0.017 |
| ***5.3.1.16: isomerase*** | 0.012 | 0.011 | 0.009 | 0.002 | 0.003 | 0.003 | 0.000 | 0.128 | 0.006 |
| ***5.3.1.24: Phosphoribosylanthranilate isomerase*** | 0.011 | 0.010 | 0.008 | 0.002 | 0.004 | 0.004 | 0.000 | 0.119 | 0.019 |
| ***5.4.99.18: 5-(carboxyamino)imidazole ribonucleotide mutase*** | 0.018 | 0.015 | 0.012 | 0.004 | 0.004 | 0.003 | 0.000 | 0.000 | 0.002 |
| ***6.1.1.22: Asparagine--tRNA ligase*** | 0.018 | 0.016 | 0.014 | 0.003 | 0.004 | 0.004 | 0.000 | 0.175 | 0.021 |
| ***6.2.1.1: Acetate--CoA ligase*** | 0.002 | 0.001 | 0.001 | 0.002 | 0.001 | 0.001 | 0.142 | 0.000 | 0.007 |
| ***6.3.2.2: Glutamate--cysteine ligase*** | 0.001 | 0.001 | 0.004 | 0.001 | 0.001 | 0.005 | 0.000 | 0.913 | 0.000 |
| ***6.3.2.5: Phosphopantothenate--cysteine ligase*** | 0.003 | 0.003 | 0.002 | 0.002 | 0.003 | 0.001 | 0.000 | 0.026 | 0.008 |
| ***6.3.4.20: 7-cyano-7-deazaguanine synthase*** | 0.009 | 0.009 | 0.007 | 0.003 | 0.004 | 0.002 | 0.000 | 0.709 | 0.000 |
| ***6.3.5.4: Asparagine synthase (glutamine-hydrolyzing)*** | 0.007 | 0.003 | 0.005 | 0.003 | 0.003 | 0.003 | 0.006 | 0.000 | 0.002 |
| ***6.4.1.2: Acetyl-CoA carboxylase*** | 0.018 | 0.011 | 0.015 | 0.005 | 0.005 | 0.006 | 0.042 | 0.000 | 0.001 |
|  | **Normal-BA group is significantly higher than High-BA group** | | | | | |  |  |  |
| * FDR: Dunn tests adjusted by Benjamini–Hochberg |  |  |  |  |  |  |  |  |  |

| **Table S9 Differential pathway among three group** | | | | | | | | | |
| --- | --- | --- | --- | --- | --- | --- | --- | --- | --- |
| **KEGG pathway** | **Health_mean** | **Normal-BA_mean** | **High-BA_mean** | **Health_sd** | **Normal-BA_sd** | **High-BA_sd** | **FDR(High-BA vs Health)*** | **FDR(Normal-BA vs Health)** | **FDR(High-BA vs Normal-BA)** |
| ko00020: Citrate cycle (TCA cycle) | 0.052 | 0.061 | 0.086 | 0.024 | 0.040 | 0.044 | 0.000 | 0.112 | 0.047 |
| ko00310: Lysine degradation | 0.010 | 0.012 | 0.028 | 0.004 | 0.004 | 0.027 | 0.000 | 0.070 | 0.045 |
| ko00362: Benzoate degradation | 0.009 | 0.008 | 0.015 | 0.002 | 0.004 | 0.013 | 1.000 | 0.003 | 0.002 |
| ko00531: Glycosaminoglycan degradation | 0.034 | 0.063 | 0.039 | 0.029 | 0.045 | 0.026 | 0.435 | 0.002 | 0.042 |
| ko00564: Glycerophospholipid metabolism | 0.026 | 0.031 | 0.051 | 0.008 | 0.012 | 0.034 | 0.000 | 0.130 | 0.018 |
| ko00627: Aminobenzoate degradation | 0.005 | 0.005 | 0.017 | 0.003 | 0.004 | 0.019 | 0.002 | 0.448 | 0.027 |
| ko00633: Nitrotoluene degradation | 0.007 | 0.008 | 0.020 | 0.003 | 0.010 | 0.024 | 0.010 | 0.473 | 0.000 |
| ko00791: Atrazine degradation | 0.002 | 0.002 | 0.010 | 0.005 | 0.003 | 0.013 | 0.010 | 0.974 | 0.022 |
| ko00920: Sulfur metabolism | 0.027 | 0.038 | 0.064 | 0.025 | 0.039 | 0.039 | 0.000 | 0.274 | 0.003 |
| ko01053: Biosynthesis of siderophore group nonribosomal peptides | 0.004 | 0.005 | 0.018 | 0.004 | 0.003 | 0.024 | 0.000 | 0.050 | 0.033 |
| ko02030: Bacterial chemotaxis | 0.036 | 0.036 | 0.071 | 0.020 | 0.036 | 0.060 | 0.046 | 0.220 | 0.000 |
| ko02060: Phosphotransferase system (PTS) | 0.014 | 0.013 | 0.062 | 0.011 | 0.009 | 0.077 | 0.001 | 1.000 | 0.001 |
| ko04974: Protein digestion and absorption | 0.004 | 0.006 | 0.004 | 0.003 | 0.004 | 0.002 | 1.000 | 0.011 | 0.025 |
| ko05100: Bacterial invasion of epithelial cells | 0.000 | 0.000 | 0.005 | 0.001 | 0.000 | 0.009 | 0.004 | 1.000 | 0.003 |
| ko05142: Chagas disease | 0.000 | 0.000 | 0.001 | 0.000 | 0.000 | 0.001 | 0.010 | 1.000 | 0.002 |
| ko00364: Fluorobenzoate degradation | 0.000 | 0.001 | 0.011 | 0.000 | 0.002 | 0.018 | 0.000 | 0.000 | 0.001 |
|  | **Normal-BA group is significantly higher than High-BA group** | | | | | | |  |  |
| * FDR: Dunn tests adjusted by Benjamini–Hochberg | |  |  |  |  |  |  |  |  |

| **Table S10 The abundance change of BA-metabolizing MAGs between NAFLD patients and Controls in the validation cohort** | | | | | | | | | | | | | | | | |
| --- | --- | --- | --- | --- | --- | --- | --- | --- | --- | --- | --- | --- | --- | --- | --- | --- |
| **mag ID** | **Scientific name** | **BSH** | **7α-HSDH** | **baiA** | **baiB** | **baiCD** | **baiE** | **baiF** | **baiH** | **baiI** | **H_mean** | **N_mean** | **H_sd** | **N_sd** | **pvalue** | **FDR** |
| mag_01 | <T>Bacteroides_vulgatus_ATCC_8482_uid58253 | 13 | 3 | 11 | 11 | 5 | 0 | 0 | 13 | 0 | 3.3442 | 8.7703 | 2.2683 | 9.0231 | 0.8928 | 0.9673 |
| mag_02 | Faecalibacterium_prausnitzii MAG02 | 5 | 4 | 2 | 3 | 2 | 0 | 0 | 6 | 0 | 1.2848 | 4.6720 | 0.6221 | 4.5296 | 0.4818 | 0.9523 |
| mag_03 | Eubacterium_rectale MAG03 | 3 | 0 | 0 | 3 | 1 | 0 | 0 | 4 | 0 | 2.4515 | 7.4117 | 1.9134 | 7.4762 | 0.6699 | 0.9523 |
| mag_04 | <T>Prevotella_dentalis_DSM_3688_uid184818 | 4 | 0 | 6 | 3 | 0 | 0 | 0 | 1 | 0 | 15.3300 | 0.0000 | 29.3411 | 0.0000 | 0.0169 | 0.2264 |
| mag_05 | Prevotella_ruminicola MAG05 | 0 | 0 | 2 | 0 | 0 | 0 | 0 | 1 | 0 | 6.5512 | 0.0000 | 10.1699 | 0.0000 | 0.0237 | 0.2644 |
| mag_06 | Ruminococcus_bromii MAG06 | 5 | 2 | 6 | 5 | 2 | 0 | 0 | 3 | 0 | 3.2238 | 6.0057 | 4.8867 | 9.6765 | 0.9280 | 0.9673 |
| mag_07 | Bacteroides_xylanisolvens MAG07 | 1 | 0 | 4 | 1 | 3 | 0 | 0 | 4 | 0 | 1.2302 | 5.1718 | 0.5041 | 6.1841 | 0.7596 | 0.9523 |
| mag_08 | <T>Faecalibacterium_prausnitzii_L2_6_uid197183 | 4 | 0 | 2 | 4 | 1 | 0 | 2 | 3 | 0 | 1.0779 | 6.9448 | 0.1424 | 8.2969 | 0.4504 | 0.9523 |
| mag_09 | Ruminococcus_bromii MAG09 | 4 | 1 | 2 | 3 | 2 | 0 | 0 | 3 | 0 | 4.4724 | 3.3735 | 12.3449 | 6.6093 | 0.0950 | 0.6367 |
| mag_10 | <T>Bacteroides_helcogenes_P_36_108_uid62135 | 7 | 3 | 6 | 5 | 2 | 0 | 0 | 5 | 0 | 3.4262 | 10.3942 | 3.8791 | 12.5003 | 0.7243 | 0.9523 |
| mag_11 | Blautia_obeum MAG11 | 2 | 0 | 0 | 2 | 2 | 0 | 0 | 2 | 0 | 1.5968 | 6.9610 | 1.5313 | 3.2606 | 0.2606 | 0.7936 |
| mag_12 | <T>Butyrate_producing_bacterium_SSC_2_uid197181 | 4 | 0 | 0 | 3 | 3 | 0 | 1 | 5 | 0 | 1.8808 | 6.0918 | 1.9970 | 4.3876 | 0.4974 | 0.9523 |
| mag_13 | Coprococcus_sp_ART55 MAG13 | 4 | 1 | 2 | 4 | 0 | 0 | 0 | 2 | 0 | 2.4309 | 2.1937 | 3.0028 | 3.8096 | 0.1739 | 0.6853 |
| mag_14 | <T>Filifactor_alocis_ATCC_35896_uid46625 | 3 | 0 | 7 | 2 | 0 | 0 | 0 | 1 | 0 | 9.4176 | 6.1067 | 20.7514 | 19.3110 | 0.8674 | 0.9673 |
| mag_15 | T>Eubacterium_eligens_ATCC_27750_uid59171 | 9 | 0 | 1 | 9 | 1 | 0 | 0 | 4 | 0 | 4.0547 | 4.1127 | 2.8511 | 5.2723 | 0.0138 | 0.2264 |
| mag_16 | Coprococcus_sp_ART55 MAG16 | 4 | 1 | 4 | 4 | 1 | 0 | 0 | 2 | 0 | 1.3106 | 3.2000 | 2.8874 | 6.5334 | 0.8067 | 0.9673 |
| mag_17 | <T>Roseburia_hominis_A2_183_uid73419 | 2 | 1 | 0 | 2 | 2 | 0 | 0 | 2 | 0 | 4.3161 | 1.3800 | 8.8067 | 3.2650 | 0.0007 | 0.0229 |
| mag_18 | Blautia_obeum MAG18 | 2 | 0 | 2 | 2 | 4 | 0 | 0 | 6 | 0 | 1.6658 | 8.8150 | 1.0219 | 7.9123 | 0.2028 | 0.7385 |
| mag_19 | Fibrobacter_succinogenes MAG19 | 6 | 0 | 1 | 5 | 1 | 0 | 0 | 1 | 0 | 4.6464 | 1.8183 | 7.7559 | 3.2033 | 0.1397 | 0.6777 |
| mag_20 | Rickettsia MAG20 | 3 | 0 | 1 | 2 | 2 | 0 | 0 | 2 | 0 | 0.7686 | 6.8487 | 1.2273 | 14.4018 | 0.6373 | 0.9523 |
| mag_21 | T>Bacteroides_helcogenes_P_36_108_uid62135 | 0 | 3 | 2 | 0 | 0 | 0 | 0 | 1 | 0 | 2.3377 | 8.3182 | 1.4827 | 7.6899 | 0.5778 | 0.9523 |
| mag_22 | Eubacterium_rectale MAG22 | 0 | 0 | 0 | 0 | 2 | 0 | 0 | 4 | 0 | 1.7048 | 9.1440 | 1.1697 | 11.0184 | 0.5144 | 0.9523 |
| mag_23 | <T>Bacteroides_vulgatus_ATCC_8482_uid58253 | 2 | 0 | 3 | 1 | 2 | 0 | 0 | 4 | 0 | 3.1094 | 6.0835 | 1.8252 | 3.5014 | 0.8530 | 0.9673 |
| mag_24 | Ruminococcus_bromii MAG24 | 4 | 1 | 3 | 4 | 0 | 0 | 0 | 1 | 0 | 0.1836 | 3.7880 | 0.4090 | 6.6096 | 0.7675 | 0.9523 |
| mag_25 | <S>Eubacterium_eligens<T>Eubacterium_eligens_ATCC_27750_uid59171 | 2 | 1 | 1 | 2 | 0 | 0 | 0 | 1 | 0 | 6.6820 | 0.0000 | 7.8714 | 0.0000 | 0.0045 | 0.1000 |
| mag_26 | Veillonellaceae MAG26 | 0 | 0 | 1 | 0 | 0 | 0 | 0 | 0 | 0 | 0.1861 | 5.5105 | 0.6171 | 8.1443 | 0.3704 | 0.9523 |
| mag_27 | Eubacterium_rectale MAG27 | 1 | 0 | 1 | 1 | 2 | 0 | 0 | 3 | 0 | 2.0842 | 12.7910 | 1.3261 | 11.4814 | 0.1552 | 0.6777 |
| mag_28 | <T>Alistipes_finegoldii_DSM_17242_uid168180 | 4 | 0 | 1 | 3 | 0 | 0 | 0 | 1 | 0 | 0.7182 | 3.5540 | 0.9723 | 3.7357 | 0.2364 | 0.7543 |
| mag_29 | <T>Bacteroides_salanitronis_DSM_18170_uid63269 | 3 | 1 | 0 | 3 | 0 | 0 | 0 | 2 | 0 | 5.1264 | 5.2900 | 13.7006 | 16.7284 | 0.7447 | 0.9523 |
| mag_30 | Eubacterium_rectale MAG30 | 0 | 1 | 0 | 0 | 0 | 0 | 0 | 1 | 0 | 1.9552 | 4.8340 | 1.4387 | 5.6636 | 0.9529 | 0.9673 |
| mag_31 | Faecalibacterium_prausnitzii MAG31 | 2 | 0 | 2 | 2 | 0 | 0 | 0 | 1 | 0 | 1.4185 | 4.6070 | 0.6276 | 5.7953 | 0.7433 | 0.9523 |
| mag_32 | <T>Eubacterium_eligens_ATCC_27750_uid59171 | 0 | 0 | 0 | 0 | 1 | 0 | 0 | 2 | 0 | 1.3370 | 5.8067 | 0.6027 | 3.8576 | 0.2204 | 0.7385 |
| mag_33 | Coprococcus_sp_ART55 MAG33 | 1 | 0 | 1 | 1 | 0 | 0 | 0 | 0 | 0 | 2.0968 | 3.0767 | 2.7529 | 6.4914 | 0.1510 | 0.6777 |
| mag_34 | <T>Bifidobacterium_adolescentis_ATCC_15703_uid58559 | 1 | 0 | 0 | 1 | 0 | 0 | 0 | 1 | 0 | 0.7911 | 6.4422 | 0.5418 | 8.5578 | 0.0886 | 0.6367 |
| mag_35 | Acidaminococcus MAG35 | 1 | 0 | 1 | 1 | 1 | 0 | 0 | 4 | 0 | 0.7127 | 3.5395 | 2.0542 | 5.6371 | 0.3487 | 0.9523 |
| mag_36 | Roseburia_intestinalis MAG36 | 0 | 0 | 0 | 0 | 3 | 0 | 0 | 4 | 0 | 2.0379 | 3.1260 | 3.9344 | 4.0527 | 0.7134 | 0.9523 |
| mag_37 | Alistipes MAG37 | 1 | 0 | 2 | 1 | 0 | 0 | 0 | 0 | 0 | 1.9436 | 6.1253 | 2.2881 | 7.0424 | 0.6889 | 0.9523 |
| mag_38 | Faecalibacterium_prausnitzii MAG38 | 4 | 0 | 0 | 4 | 0 | 0 | 0 | 0 | 0 | 0.6115 | 4.4930 | 0.5047 | 5.6480 | 0.1618 | 0.6777 |
| mag_39 | <T>Prevotella_sp_oral_taxon_299_str_F0039_uid45899 | 0 | 0 | 2 | 0 | 0 | 0 | 0 | 0 | 0 | 3.1755 | 0.0000 | 5.1117 | 0.0000 | 0.0838 | 0.6367 |
| mag_40 | MAG40 | 0 | 0 | 0 | 0 | 0 | 0 | 0 | 0 | 0 | 4.3573 | 12.6402 | 7.3833 | 12.4956 | 0.6102 | 0.9523 |
| mag_41 | <T>Bacteroides_vulgatus_ATCC_8482_uid58253 | 1 | 1 | 1 | 1 | 0 | 0 | 0 | 3 | 0 | 2.6583 | 6.3073 | 1.9597 | 8.2041 | 0.9304 | 0.9673 |
| mag_42 | <T>Ruminococcus_champanellensis_18P13_uid197169 | 2 | 1 | 1 | 2 | 1 | 0 | 0 | 1 | 0 | 0.8476 | 3.4433 | 2.1399 | 7.5562 | 0.2176 | 0.7385 |
| mag_43 | <T>Bacteroides_vulgatus_ATCC_8482_uid58253. | 0 | 1 | 0 | 0 | 0 | 0 | 0 | 0 | 0 | 2.6671 | 5.7395 | 2.4497 | 8.5491 | 0.8242 | 0.9673 |
| mag_44 | <T>Bacteroides_vulgatus_ATCC_8482_uid58253. | 1 | 0 | 0 | 1 | 0 | 0 | 0 | 0 | 0 | 5.1930 | 10.2430 | 3.5932 | 4.5828 | 0.7072 | 0.9523 |
| mag_45 | Faecalibacterium MAG45 | 2 | 0 | 1 | 2 | 0 | 0 | 0 | 1 | 0 | 1.3823 | 5.8078 | 0.3858 | 6.7983 | 0.5747 | 0.9523 |
| mag_46 |  | 0 | 0 | 0 | 0 | 0 | 0 | 0 | 1 | 0 | 2.8442 | 7.8377 | 2.5756 | 10.2580 | 0.9420 | 0.9673 |
| mag_47 | Faecalibacterium_prausnitzii MAG47 | 1 | 0 | 0 | 1 | 0 | 0 | 0 | 0 | 0 | 1.2458 | 5.8280 | 0.2174 | 6.9852 | 0.5551 | 0.9523 |
| mag_48 | Coprococcus_sp_ART55 MAG48 | 1 | 0 | 2 | 1 | 0 | 0 | 0 | 0 | 0 | 1.9470 | 1.7043 | 3.1214 | 5.0117 | 0.0320 | 0.3067 |
| mag_49 | MAG49 | 0 | 0 | 0 | 0 | 0 | 0 | 0 | 0 | 0 | 3.3824 | 0.6913 | 3.8433 | 0.6087 | 0.0003 | 0.0203 |
| mag_50 | Bacteroides MAG50 | 1 | 0 | 1 | 0 | 0 | 0 | 0 | 1 | 0 | 1.9285 | 2.2512 | 3.6801 | 3.7065 | 0.1275 | 0.6777 |
| mag_51 | Faecalibacterium MAG51 | 1 | 0 | 0 | 1 | 0 | 0 | 0 | 0 | 0 | 1.1555 | 4.2260 | 0.5261 | 5.1589 | 0.6191 | 0.9523 |
| mag_52 | <T>Bacteroides_helcogenes_P_36_108_uid62135 | 1 | 0 | 0 | 0 | 0 | 0 | 0 | 2 | 0 | 2.6115 | 5.9732 | 2.9652 | 8.7108 | 0.9241 | 0.9673 |
| mag_53 | <T>Filifactor_alocis_ATCC_35896_uid46625 | 1 | 0 | 1 | 1 | 0 | 0 | 3 | 0 | 0 | 1.6688 | 2.7262 | 1.9365 | 4.4750 | 0.1239 | 0.6777 |
| mag_54 | <T>Ruminococcus_albus_7_uid51721 | 1 | 0 | 0 | 1 | 0 | 0 | 0 | 0 | 0 | 2.2039 | 4.8227 | 2.9161 | 6.8995 | 0.9685 | 0.9685 |
| mag_55 | Prevotella_ruminicola MAG55 | 0 | 0 | 1 | 0 | 1 | 0 | 0 | 1 | 0 | 1.4659 | 1.5750 | 3.2641 | 4.9806 | 0.5485 | 0.9523 |
| mag_56 | Bacteroides_xylanisolvens MAG56 | 0 | 1 | 0 | 0 | 1 | 0 | 0 | 1 | 0 | 1.1255 | 5.5065 | 0.4351 | 6.2220 | 0.5483 | 0.9523 |
| mag_57 | <T>Clostridium_sp_SY8519_uid68705 | 0 | 0 | 0 | 0 | 0 | 0 | 0 | 0 | 0 | 1.2503 | 4.2110 | 0.2781 | 3.6361 | 0.6311 | 0.9523 |
| mag_58 | Eubacterium_rectale MAG58 | 0 | 0 | 1 | 0 | 0 | 0 | 0 | 0 | 0 | 1.5939 | 8.8880 | 1.4110 | 12.4862 | 0.6698 | 0.9523 |
| mag_59 | MAG59 | 0 | 0 | 0 | 0 | 0 | 0 | 0 | 0 | 0 | 5.2058 | 6.6013 | 10.2949 | 9.1099 | 0.9159 | 0.9673 |
| mag_60 | Fibrobacter_succinogenes MAG60 | 0 | 0 | 0 | 0 | 0 | 0 | 0 | 0 | 0 | 1.4541 | 6.3278 | 0.7432 | 5.6776 | 0.3750 | 0.9523 |
| mag_61 | MAG61 | 0 | 0 | 1 | 0 | 0 | 0 | 0 | 0 | 0 | 2.2124 | 5.3080 | 2.0099 | 8.6107 | 0.6730 | 0.9523 |
| mag_62 | Faecalibacterium_prausnitzii MAG62 | 0 | 0 | 1 | 0 | 0 | 0 | 0 | 1 | 0 | 1.9042 | 6.9093 | 1.1585 | 9.0765 | 0.7464 | 0.9523 |
| mag_63 | Faecalibacterium_prausnitzii MAG63 | 0 | 0 | 0 | 0 | 0 | 0 | 0 | 0 | 0 | 2.8435 | 9.8200 | 0.8795 | 6.9197 | 0.3833 | 0.9523 |
| mag_64 | <T>Prevotella_sp_oral_taxon_299_str_F0039_uid4589 | 0 | 0 | 0 | 0 | 0 | 0 | 0 | 0 | 0 | 2.3636 | 16.7040 | 3.7730 | 26.7332 | 0.5296 | 0.9523 |
| mag_65 | <T>Bacteroides_thetaiotaomicron_VPI_5482_uid62913 | 0 | 0 | 0 | 0 | 0 | 0 | 0 | 0 | 0 | 2.6641 | 5.0083 | 3.6698 | 10.4203 | 0.5295 | 0.9523 |
| mag_66 | <T>Alistipes_finegoldii_DSM_17242_uid168180 | 0 | 0 | 0 | 0 | 0 | 0 | 0 | 0 | 0 | 2.2711 | 8.3798 | 2.2699 | 8.8499 | 0.4571 | 0.9523 |
| mag_67 | <T>Fluviicola_taffensis_DSM_16823_uid65271 | 0 | 0 | 0 | 0 | 0 | 0 | 0 | 0 | 0 | 2.4809 | 4.6535 | 5.0952 | 13.0113 | 0.8159 | 0.9673 |
|  | BA-metabolizing MAGs |  |  |  |  |  |  |  |  |  |  |  |  |  |  |  |
